# Analyzing the Perceptual Attributes of Electro-tactile Stimuli as Function of Various Signal Properties

**DOI:** 10.1101/2021.08.08.455590

**Authors:** Mehdi Rahimi, Fang Jiang, Yantao Shen

## Abstract

This study is designed to determine the proper electrical signals that induce various perceptual qualities. These perceptions are chosen from a wide range of feelings, comfort and even surface qualities such as smoothness or flatness. The goal is to derive the relationship between the electrical signal properties (i.e. voltage, frequency and duty cycle) and the perceived qualities which are understood by human observers and reveal any pattens in their reports. A total of 144 measurements are collected from 8 participants. The results are analyzed using numerous statistical methods. In the end, the governing factors are identified and a signal with specific properties is is determined for each of the perception qualities. A great number of plots are included in this study to facilitate the process of choosing the appropriate signal for each factor.

## I. Introduction

**A** comparison between mechanical stimulation of the skin and electro-tactile stimulation can result in showing various benefits for the electrical approach [1]. One significant factor would be the undeniable shorter time that an electrical signal needs for any changes, whereas a mechanical approach is more limited by the constraints of the moving parts of the motors and other segments of the system. [2]. The skin reaction to electrical stimulation is also considerably faster and more accurate [3][4]. Electrical stimulation also has a higher convenience level, better efficiency, and more flexibility for the purpose of giving sensory information along with less noise and lighter weight of the system [5].

Despite these and many other advantages, there are some limitations associated with the electrical stimulation. One limitation to consider, is the location of the contacts that are placed on the skin [6]. Another factor is the size and the shape of the electrodes [7]. Several studies have focused on using the anode and cathode method and placing the return contacts in a close vicinity of each other [8]. This was mainly done to stimulate different nerves for the purpose of inducing various feelings such as pressure or vibration [9]. The main problem with this approach is that putting the return electrode in a close proximity of the anode or cathode electrodes would potentially limit the area that can be used for building the electro-tactile display.

To overcome this issue, in this study, we chose to position the contacts slightly distal to the fingertip vortex and have a single-point return contact on the palm of the hand. This decision is to facilitate the construction of bigger displays on the skin in future studies. This approach has been tried before and can give an option to stimulate the Merkel cells in the skin [10][11]. The size, shape, and the material of the contacts can also play a role in sensations [7] [12]. In this study, all the contacts are copper circles with the diameter of 1.5*mm*. This can result in stimulating the receptors directly [13] it will also adhere to the two-point discrimination threshold (TPDT) of the skin in that area [14].

Another topic to consider about the electrical signal is the choice of the signal parameters (i.e. voltage, frequency and duty cycle) [15]. Several studies have focused on finding the perfect combination of signal properties to achieve an acceptable feeling of the electro-tactile stimulation [12] [11] [16]. While it is not possible to give a thorough review of all these works and many others here, the contributions of some of them are discussed. A few reviews are published by Kaczmarek and Bach-y-Rita [4] [1] that show comparisons between these works that are done in this area. Some more recent studies such as the one done by Ara et al [16] investigated the effect of 20Hz and 200Hz signals on sensations such as Beating, Fluttering, Vibrating, and Buzzing. The effect of increasing the voltage was also looked at and even though the studied frequencies are different between that work and this study, the results in this area are very similar. The work of Tahiso and Higashiyama [12] is limited to a frequency of 60Hz but their approach involved giving a series of pulses instead of a continuous signal. Their work focused on the effects of electrical current on various sensations such as Touch, Prick, Deep Pressure, and Itch. Szeto [17] found that frequencies below 100Hz are the most useful range for sensory communications. Kantor [18] investigated five waveforms with various pulse per second (PPS) rates and concluded that all of them were successful in exciting the nerve fibers in the skin. They studied a short range of variation in peak voltage. None of the studies investigated the effect of changing the duty cycle on various sensations. These and other studies show that there is not a thorough investigation of the relationship between various signal properties (voltages, frequencies and duty cycle) and different sensations. This is examined systemically in this study.

This paper is structured into the following sections: In Section II, we describe the methodology and experiments used in this study. Section III covers the results obtained from the analysis. A discussion about the results is presented in Section IV and in Section V we conclude the study.

## II. Methods

### A. Hardware and Systems

The contacts on the finger were designed as copper circles with a diameter of 1.5*mm*. They were built using a printed circuit board (PCB). The return electrode on the palm of the hand was a copper circle with a diameter of 3*cm*. A switching board was designed to amplify the input signal from a function generator (UNi AFG-1010). These were connected to a Raspberry Pi 3 with a display to show visually when and if a contact is on. Fig. 1 shows this setup. An overall sketch of the switching board is shown in Fig. 2. A high voltage DC power supply (TTi PLH250) was used to power the switching board. The current on the power supply was limited to 0.2*mA* to prevent any accidental increases in current. Furthermore, a 2*mA* fuse was put in series with the circuit as an extra precaution. An ammeter (MASTECH MS217) was also used in series to measure the current that goes through the skin. The power supply was able to generate any DC voltage from 0 to 250 Volts; although our experiments were limited to a range of 40 – 120 Volts depending upon the individual’s skin condition and sensation threshold. The function generator was capable of generating any frequencies with an accuracy of 0.0001*Hz* and any duty cycles with an accuracy of 1%. The power supply had an accuracy of 0.1*V*. Apart from all the precautionary procedures, the placement of the contacts was so that the participants could easily lift their fingers from the contacts and disconnect the circuit in case of any discomforts. Participants were also instructed to verbally communicate any discomfort or abnormal feelings.

**Figure 1.**
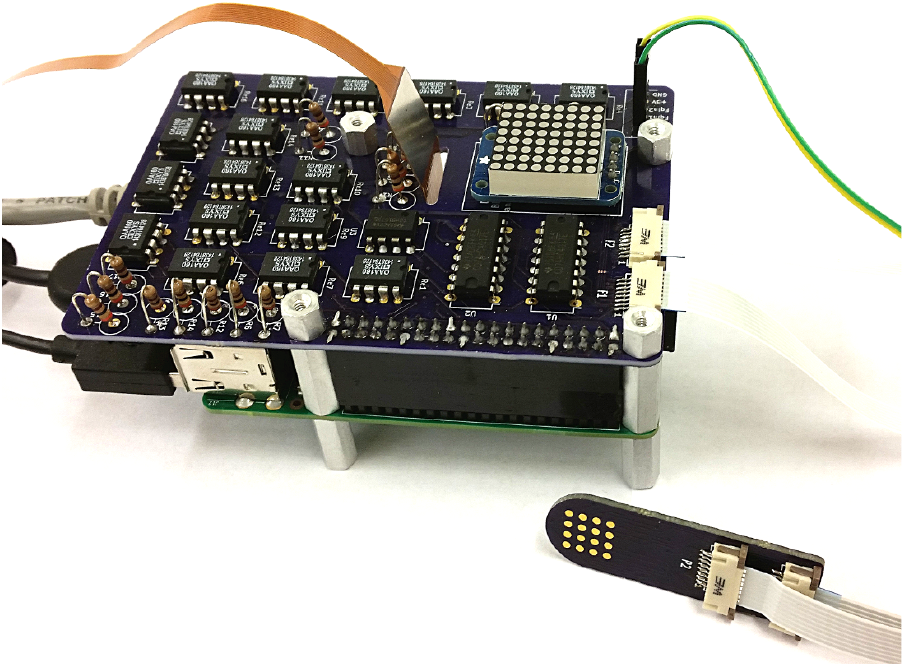
The switching board with the display are connected to the top of a Raspberry Pi. The board controls the contacts. Multiple contacts were designed to stimulate different parts of the skin.

**Figure 2.**
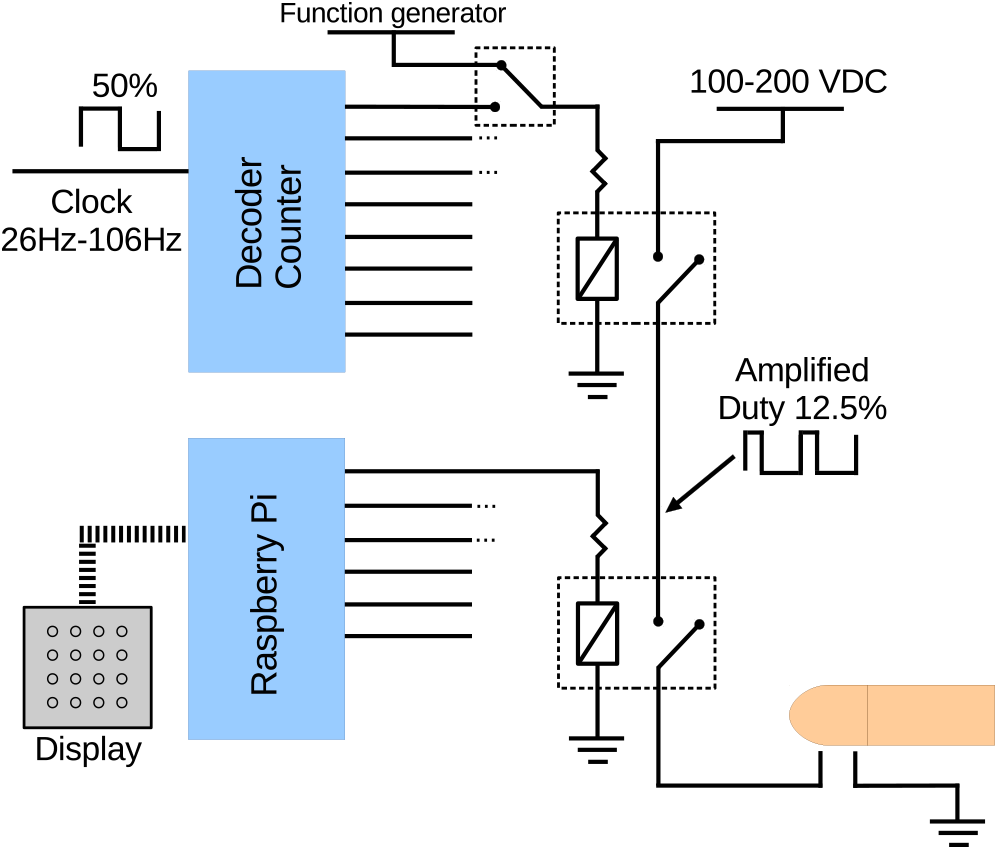
An overall sketch of the switching board is shown here. A counter can distribute the signal to regulate the duty cycle. A switch can decide between the counter or the function generator. A Raspberry Pi program controls each pin to be on or off. Relays are used to switch on the signals.

### B. Participants

In this study, eight participants (seven males and one female) took part. They were all healthy graduate students at the University of Nevada, Reno. None of them reported any physical problems nor had any issues with their skin or fingers. Participation was with informed consent and followed protocols approved by the University of Nevada, Reno Institutional Review Board. The purpose and procedure of the test were explained to each participant before taking the test. The questionnaire was also explained extensively to make sure there would not be any misunderstanding or misinterpretations. A cardboard poster-holder was placed between the participants and the system so that they could not see the voltage, frequency, or duty cycle for each experiment. This was done to prevent any prejudgements toward higher or lower values of signal properties.

### C. Signal Properties

The goal of this study was to find the proper properties for the electrical signal to stimulate the skin. Although there can be an infinite number of combinations for the signal, in this study, 18 different combinations were selected based on previous preliminary experiments. The frequencies that were tested were 10*Hz*, 30*Hz*, and 60*Hz*. Since all previous experiments were done using 30*Hz* signal, this was selected as the appropriate frequency. The duty cycles were 5% and 10%. Three voltage selections were chosen. The first was the detection threshold (DT) voltage that is identified in this study as “*V* 1”. This was identified using a staircase method [19]. The two other voltages were defined as *V* 2 = *V* 1 + 10%(*V* 1) and *V* 3 = *V* 1 + 20%(*V* 1). These selections resulted in 18 combinations that were tested on each participant.

### D. Questionnaire

To assess the perceived qualities of each electrical signals, a questionnaire was designed based on previous research in this area [16] [20] [12]. The questionnaire had two parts.

In the first part, the participant would choose one or more of the feelings with these explanations provided: Nothing, Slight tough, Pressure (as if an object is pushing the skin), Deep Pressure (as if the push is intense and can be felt deep inside the tissue), Vibrating (beating, pulsation, throbbing), Itch (the urge to scratch the skin), Sting (prick, sharp pain like a needle), Unpleasant (when the participant simply did not like the sensation or the signal was causing any pain). The second part was a more quantitative measure where 7 feelings were shown on a scale of 0-10 and the participant had to choose a number in that range. It was explained to them that selecting 5 means neutral feeling. The 7 feelings and their explanations were: Comfortable vs. Uncomfortable, Low-frequency vibration vs. High-frequency vibration (the feeling of the signal on the skin), Surface and Shallow sensation vs. Deep Sensation (the feeling that the sensation in on the surface of the skin versus when it is felt deep inside the tissue), Smooth vs. Rough (as an example of leather vs. sand paper), Flat vs. Bumpy (as an example of tile vs. sponge), Hard surface vs. Soft surface (as an example of brick vs. velvet) and finally, Warm vs. Cold (reports of any temperature feelings on the skin). The questionnaire is shown in Fig. 3.

**Figure 3.**
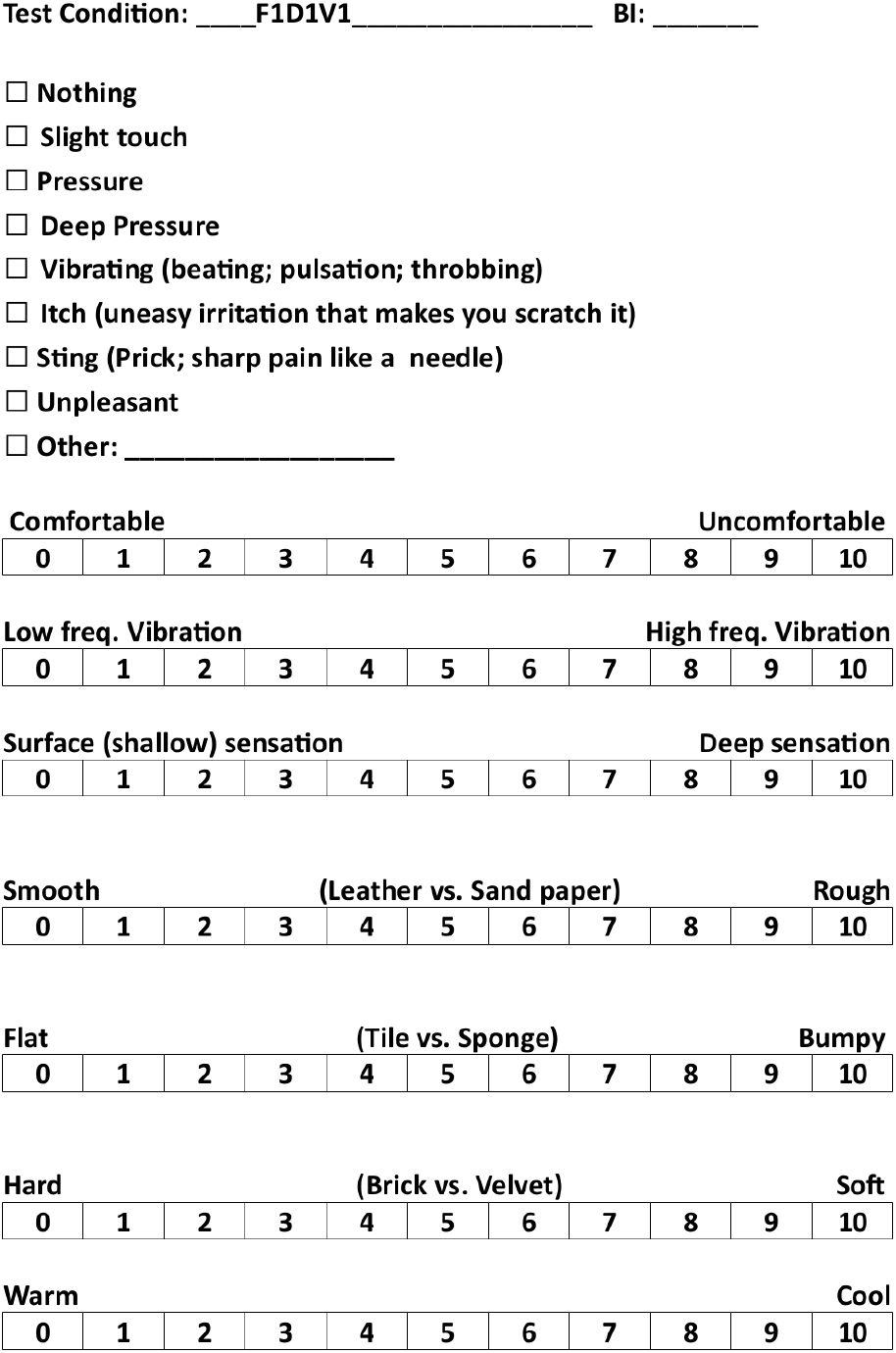
The questionnaire used in this study is shown here. This was filled by the participants after each and every experiments.

### E. Experimental Procedure

To prepare for each set of experiments, an examination of each part of the system was conducted to ensure every part of the system was performing as expected. Then, the participant was briefed on the experiment and what they needed to do in each part. A thorough explanation of the questionnaire was also included to mitigate any misunderstandings. The contacts and the participant’s finger and palm were then cleaned using Ethyl Alcohol 200 Proof and left to air dry. The subjects could not see the system as there was a cardboard poster-holder placed between them and the system to prevent any prejudgements about the signal properties. Each experiment was started by identifying the detection threshold (DT) voltage (i.e V1). The subjects were given a time between 5-20 seconds to examine the feeling and then were instructed to lift their fingers and therefore disconnect the circuit. The subject then started to fill out the questionnaire until they were ready for the next experiment.

A phenomenon called skin tissue fatigue occurs when the skin is exposed to the electrical signal for a duration of time. This can decrease the sensitivity of the skin and interfere with the feelings perceived. To overcome this, the experiments were conducted in two sessions. Since the whole test for each individual could take about 40 minutes, a 30-minute rest was used after 20 minutes of the experiment. The subject could use this rest time to leave the station and come back after that. Cleaning of the contacts, finger, and palm was repeated after this rest and before continuing the experiment.

The voltage and current for each experiment were recorded to calculate V2 and V3 and also making sure the placement of the finger was identical to the previous test. These data along with the test conditions were written on the questionnaire in coded format to prevent the psychological side effects on the participants.

## III. Results

### A. Results from Part I of the questionnaire

As a result of the extensive questionnaire used in this study, various analysis were performed on the data. We first examined how the subjects perceived all these test conditions. Fig. 4 shows the overall perception of all the various test signals.

**Figure 4.**
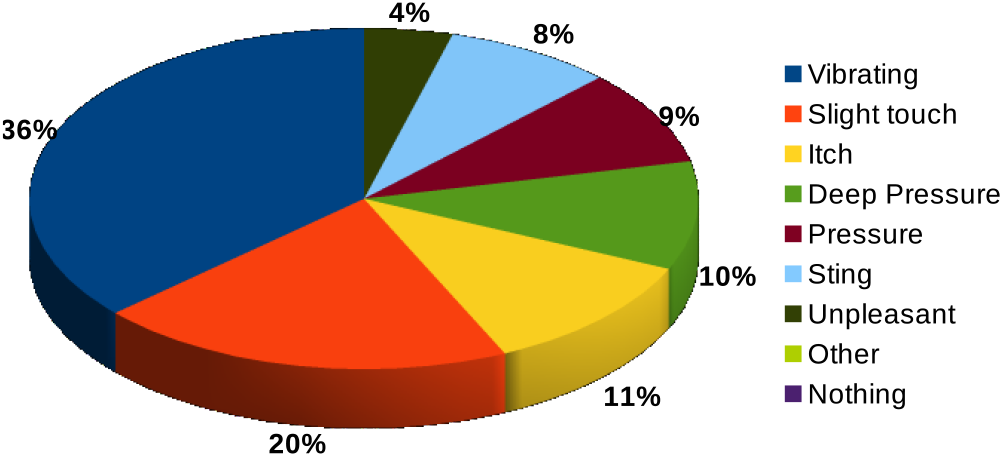
The participants described each signal and the overall assessment is shown here. The most dominant sensation is vibrating, as 36% of reports were that the electrical signal gives a vibrating sensation.

The fact that vibrating sensation is the most dominant across all the reported sensations illustrates an important characteristic of how subjects faced the electrical signals in general. This can also be investigated more by looking at how each test signal is perceived. Fig. 5 shows a per-experiment analysis.

**Figure 5.**
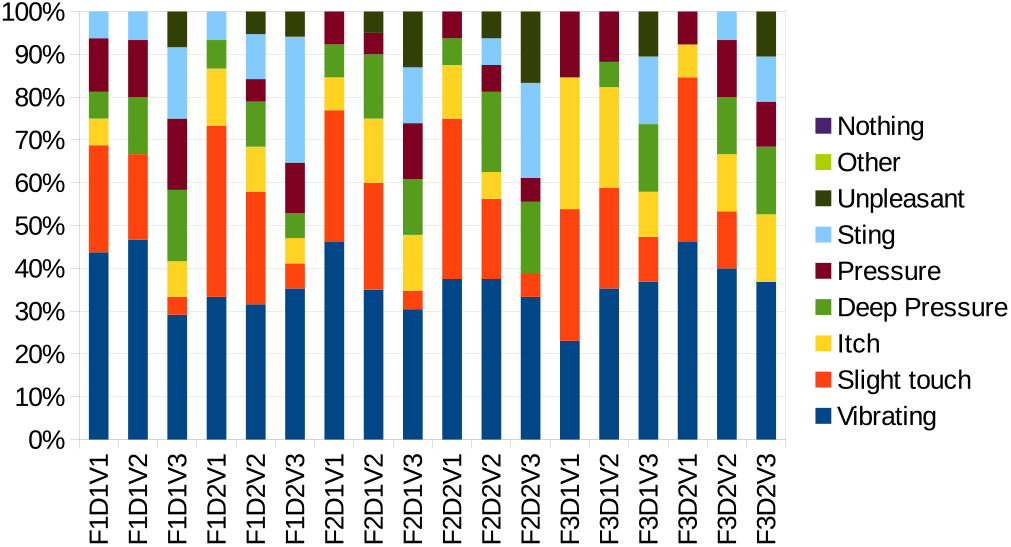
The reporting for each of the 18 test conditions is shown here. Each column is normalized to show how much of the overall assessment was contributed to each feeling. In this figure, F1, F2 and F3 represent 10Hz, 30Hz and 60Hz respectively and D1, D2 are 5% and 10% duty cycles. “*V* 1” is the detection threshold (DT) voltage. The two other voltages were defined as *V* 2 = *V* 1 + 10%(*V* 1) and *V* 3 = *V* 1 + 20%(*V* 1).

Fig. 5 shows a thorough examination of each test, but, to extract the patterns that are embedded in this figure, an area plot was used to emphasize the parameters that are harder to notice which is shown in Fig. 6.

**Figure 6.**
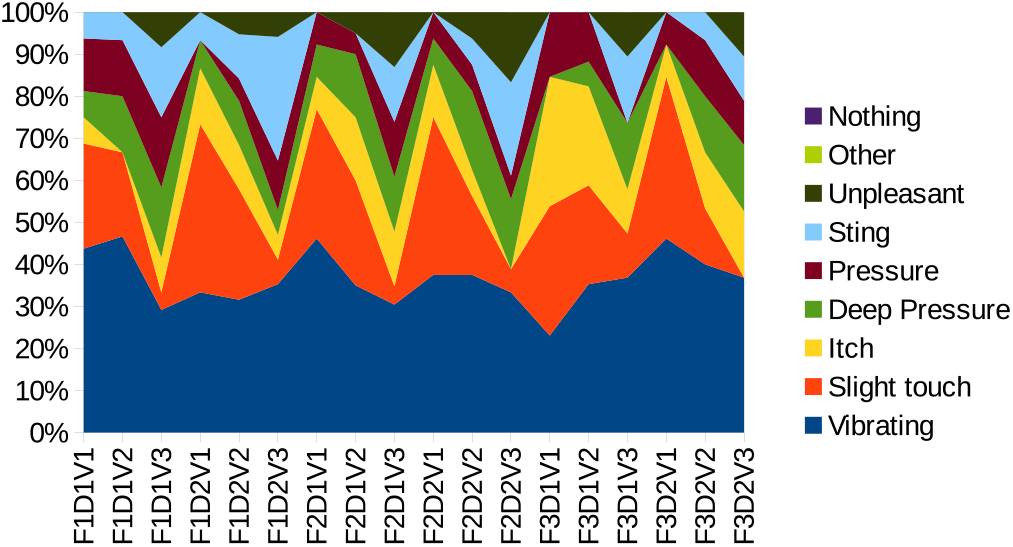
A box plot is used to show the embedded patterns of the reports across all the tests. It can be easily seen that “Slight touch”, “Sting” and “Unpleasant” are the governing factors and depend on voltage more than anything else. In this figure too, F1, F2 and F3 represent 10Hz, 30Hz and 60Hz respectively and D1, D2 are 5% and 10% duty cycles.

Fig. 6 shows that there is a definite relationship between voltage and feelings of “Slight touch”, “Sting” and “Unpleasant”. To summarize the conclusion here, a simplified version of Fig. 6 that is focused on these three reports is generated and shown in Fig. 7

**Figure 7.**
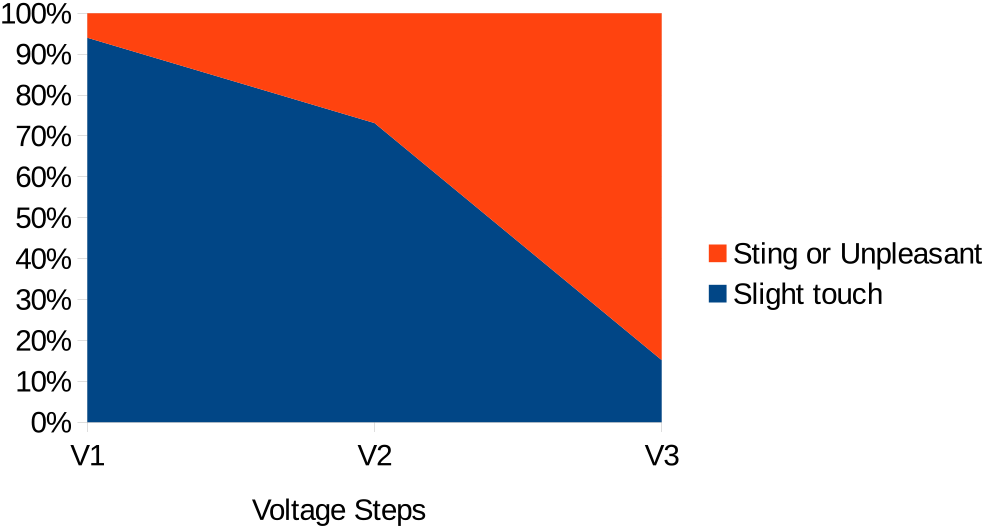
A simplified box plot of Fig. 6 is used to show the direct relationship between “Slight touch”, “Sting” and “Unpleasant” and the voltage. As discussed before, “*V* 1” is the detection threshold (DT) voltage. The two other voltages were defined as *V* 2 = *V* 1 + 10%(*V* 1) and *V* 3 = *V* 1 + 20%(*V* 1).

Fig. 7 is summarizing Fig. 6 in the most concise way possible. It is shown that in lower voltages, the dominant reports from the participants were “Slight touch” but as we increased the voltage pass the DT point, that feeling was converted to either “Sting” or simply “Unpleasant”. We will see a similar pattern later in this study and it will be discussed furthermore.

We can extend these analyzes to see how these sensations reports are actually changing in relation to changes in voltage, frequency, or duty cycle. This might better reinforce the previous conclusion that voltage is the governing factor.

Fig. 8 is to show how many reports were made for each feeling and this is plotted in relation to the changes in frequency. This is done across 144 experiments that were done in this study.

**Figure 8.**
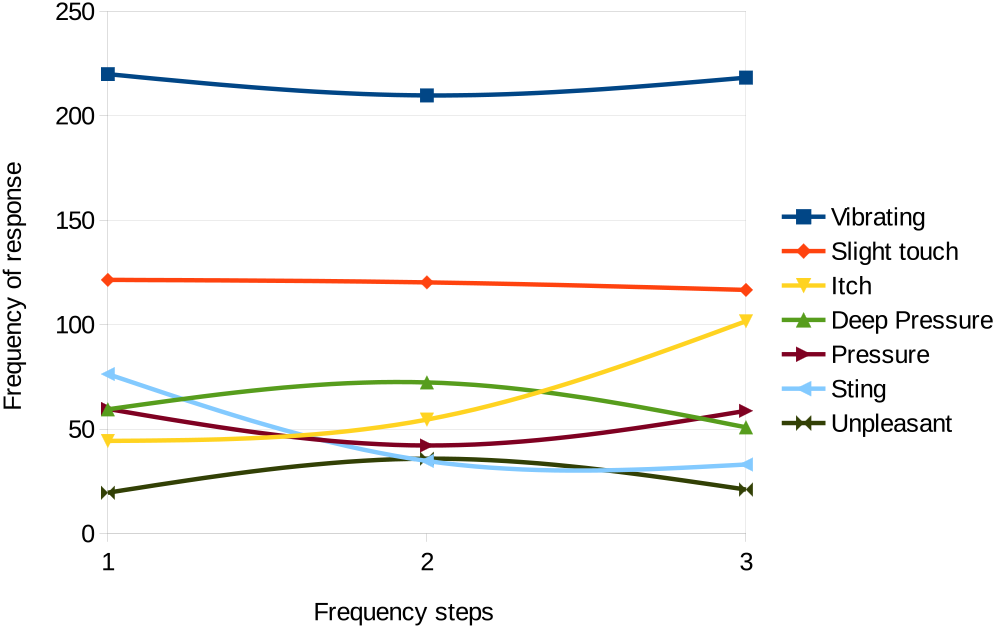
The number of times that each feeling was reported is plotted versus the different frequencies of the signals. As was shown before, “Vibrating” is the most dominant factor. But more importantly, this plot shows that most of the reports of the feeling did not change as a function of frequency. Steps of frequencies here are 10*Hz*, 30*Hz* and 60*Hz*.

One observation from Fig. 8 can be that although the reports of most of the feelings stayed the same in relation to the changes in frequency, but the “Itch” feeling did have an increase to as much as twice its initial reported value going from the lowest frequency (10*Hz*) to the highest one (60*Hz*).

We can examine the same principle considering the duty cycle. Fig. 9 is examining the relationship between the number of times that each feeling was reported as a function of changing the duty cycle.

**Figure 9.**
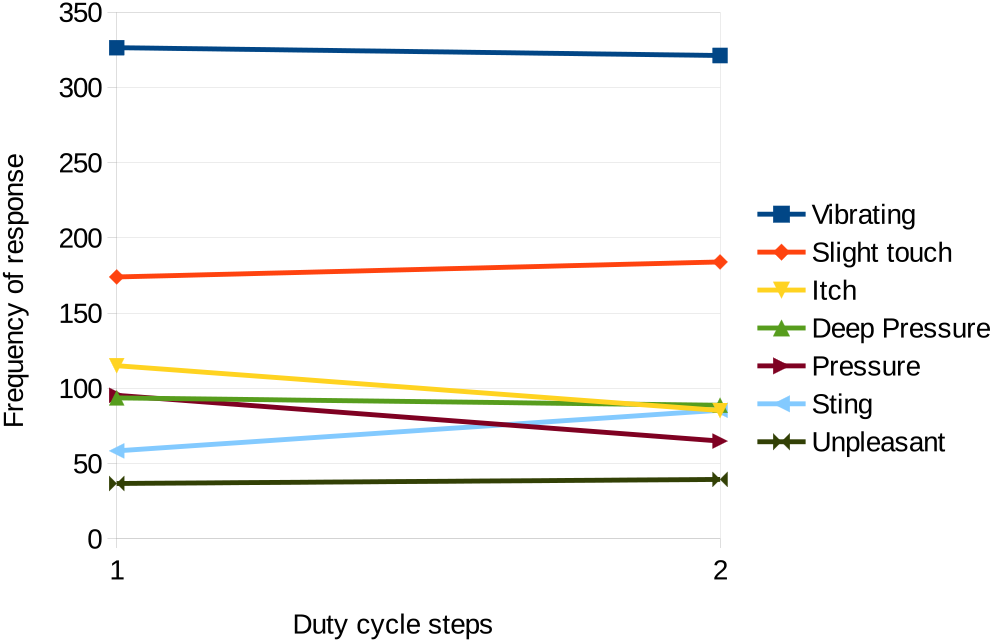
The number of times that each feeling was reported is plotted versus the different duty cycles of the signals. No significant relationship is indicated here. Steps of duty cycles here are 5% and 10%.

Fig. 9 shows that the relation between the reports and the duty cycle is even weaker than the frequency as we examined in Fig. 8. There are no significant changes in relation to the duty cycle for the number of times that each feeling was reported.

As the final step in this part, we can examine the same effect considering the changes in voltage. Fig. 10 is investigating this relationship.

**Figure 10.**
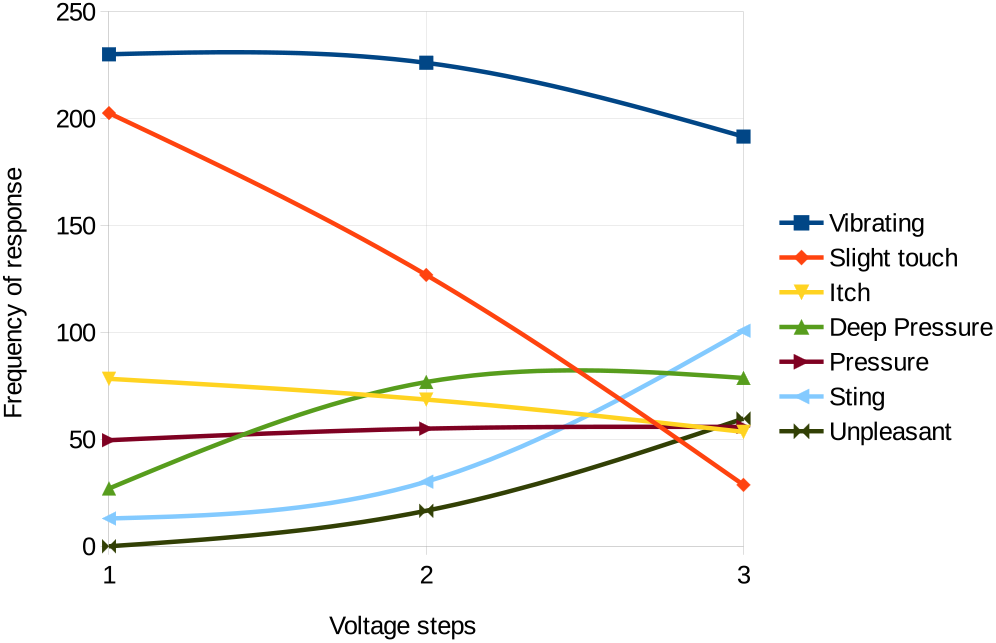
The number of times that each feeling was reported is plotted versus the different voltages of the signals. Significant changes can be seen especially in the feeling of “Slight touch”. Steps of voltages here are “*V* 1” as the detection threshold (DT) voltage, *V* 2 = *V* 1 + 10%(*V* 1) and *V* 3 = *V* 1 + 20%(*V* 1)

As much as we did not see any significant relationship between the feelings and the frequency or the duty cycle, the same cannot be said about the voltage. Fig. 10 shows that the feeling of “Slight tough” was considerably affected by increasing the voltage. The “Vibrating” feeling did show a slight decrease too, but more importantly, the feelings of “Sting” and “Unpleasant” show substantial growth with increasing the voltage. These are all further verifying our conclusions from Fig. 6 and 7.

### B. Results from Part II of the questionnaire

In the second part of the questionnaire, 7 feelings were assessed on a rating scale between 0 to 10. It was explained to the participants that choosing number 5 means “neutral”. We can analyze the relationship between each of these 7 feelings with respect to signal properties.

#### 1) Comfortable vs. Uncomfortable

The effect of changes of frequency on the feeling of comfort or discomfort can be analyzed. Fig. 11 shows this relationship. There is no significant difference between the groups meaning that there is not a notable association between the frequency and the sense of comfort.

**Figure 11.**
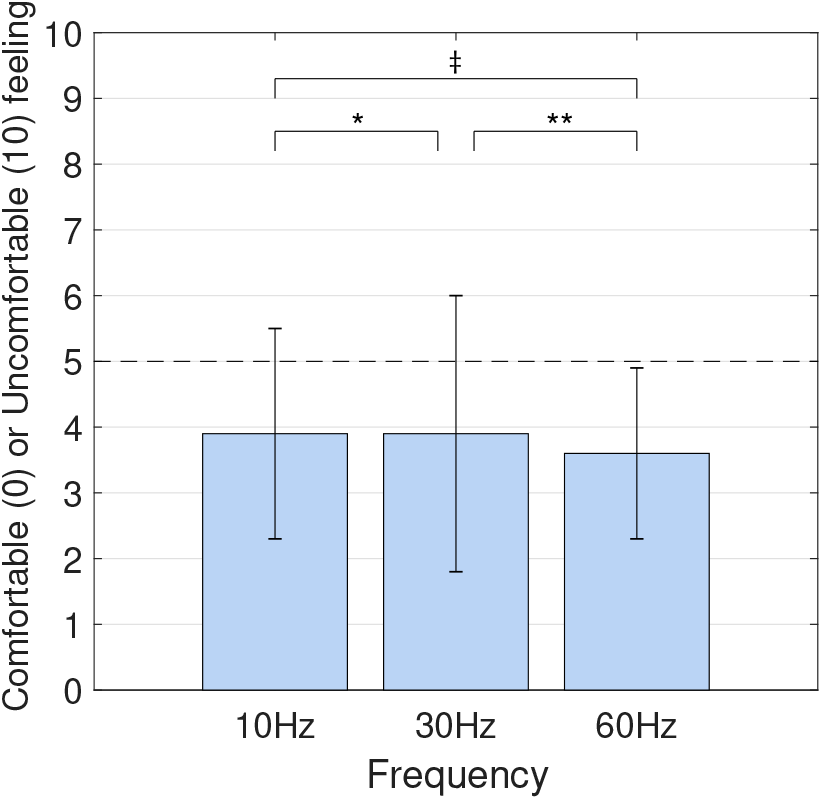
The effect of increasing frequency on the comfort or discomfort feeling is shown here. Significance of the overall differences between all groups: 0.930, ∗*p* = 0.988, ∗ ∗ *p* = 0.760, ‡*p* = 0.707

In the same way, the effect of duty cycle can be investigated too. Fig. 12 shows that there is a minute increase in discomfort when the duty cycle was increased. The difference is still not significant and is not considerable enough to be seen as a major controlling factor.

**Figure 12.**
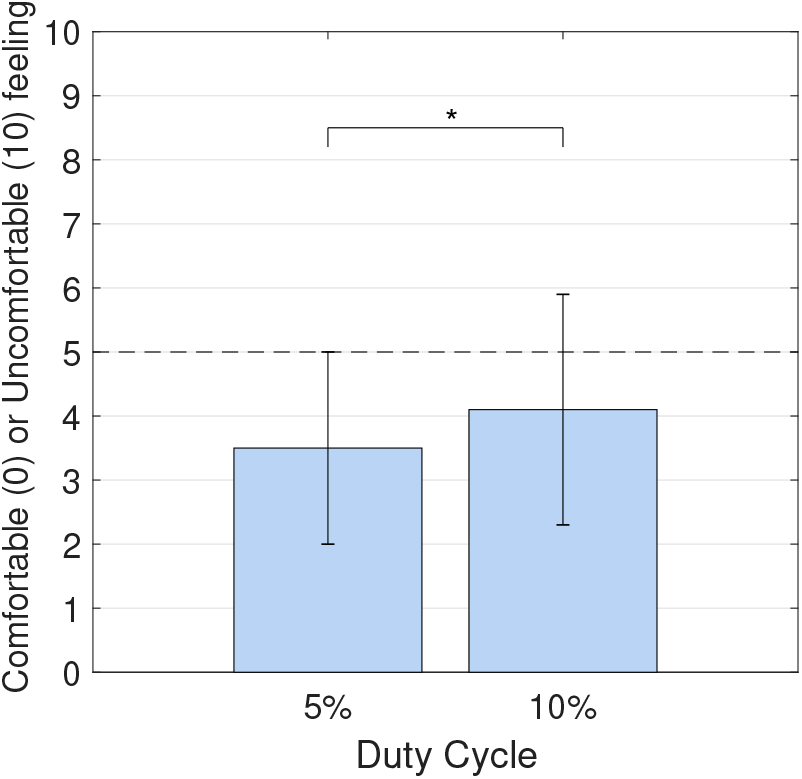
The effect of increasing duty cycle on comfort or discomfort feeling is shown here. ∗*p* = 0.481

As we expected, based on previous analysis, the voltage should have a much more important effect in this area. Fig. 13 shows this connection between the two. In this figure, we can see a noteworthy association. The overall difference groups between and also the difference between each two groups are all statistically significant (*p* < 0.001). *V*1 is clearly showing a comfortable feeling whereas increasing the voltage is changing this to the “uncomfortable” side.

**Figure 13.**
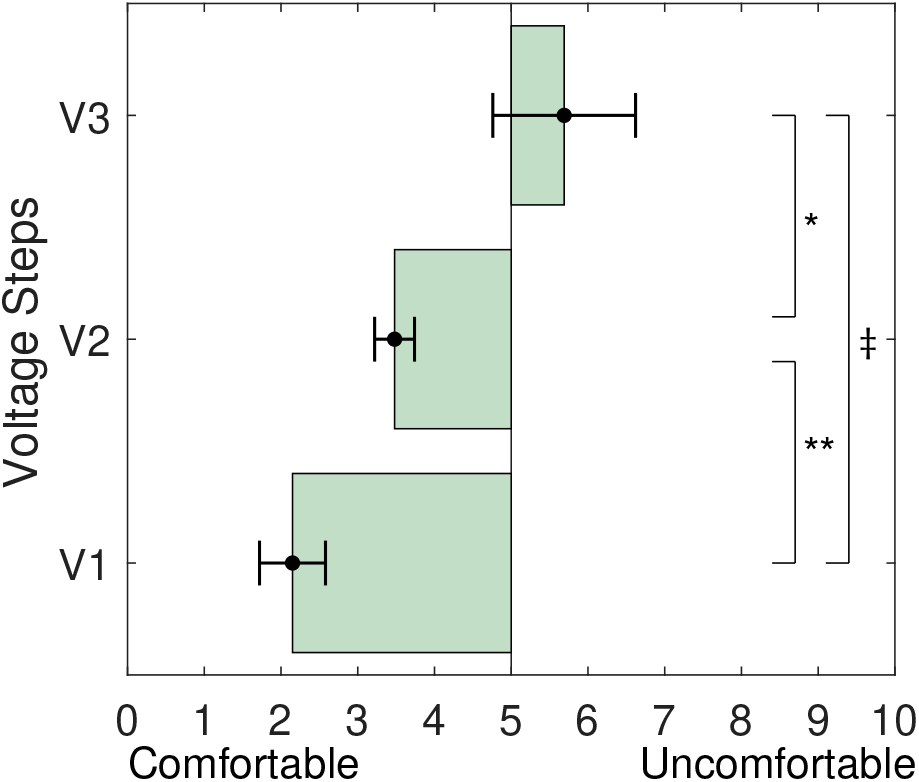
The effect of increasing voltage on comfort or discomfort feeling is shown here. Significance of the overall differences between all groups: < 0.0001, ∗*p* < 0.001, ∗ ∗ *p* < 0.0001, ‡*p* < 0.0001

We can also sort all the various test signals in connection to the comfort to find the most appropriate properties for a comfortable signal. Fig. 14 illustrates this arrangement. Noting this figure, an undeniable grouping based on the voltage rather than any other factors can be seen. We can also conclude that the “V1” voltage (DT) along with the lowest duty cycle can result in the most comfortable feeling.

**Figure 14.**
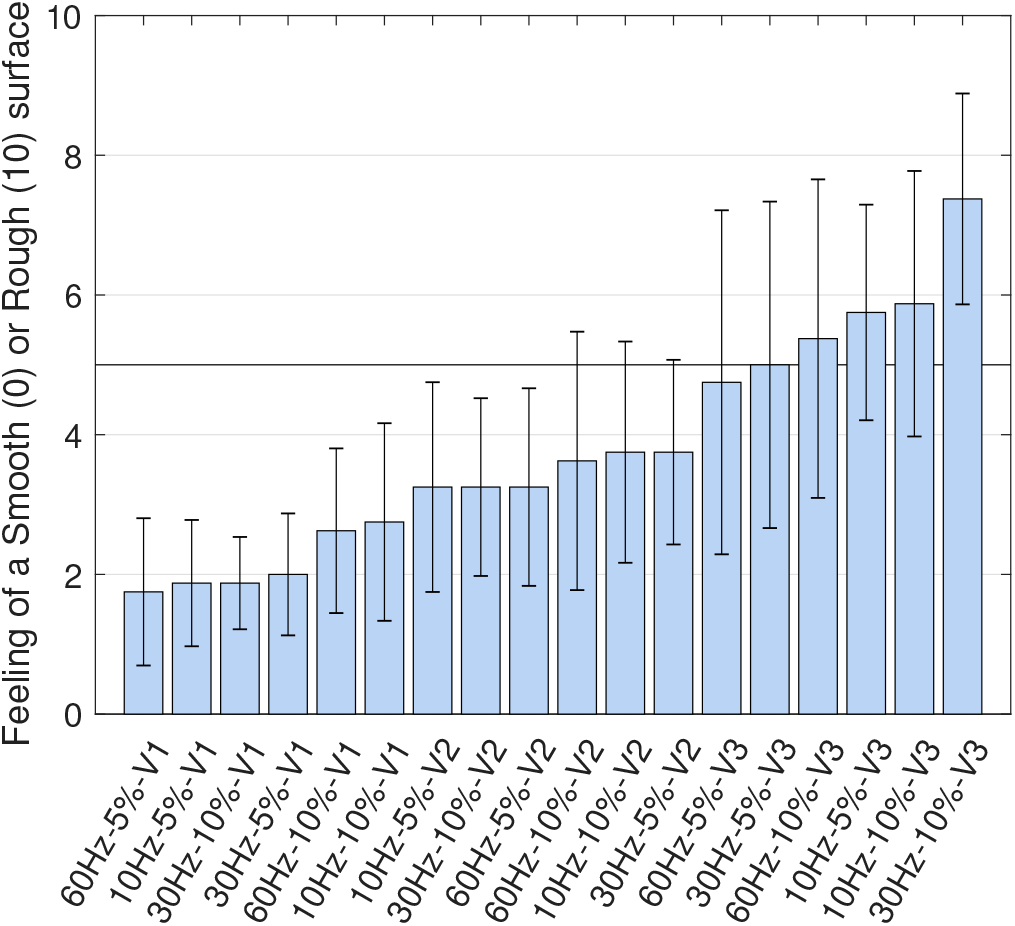
All the 18 different signals are sorted here based on the comfortable or uncomfortable feelings reports.

#### 2) Low-frequency vibration vs. High-frequency vibration

Since most participants felt the electrical stimulation as a vibration feeling on their skin, on the questionnaire we asked them about the quality of this feeling; weather if this vibration feel like a low-frequency vibration or a high-frequency. We could then analyze how different signal properties affected this feeling. Fig. 15 shows this relationship.

**Figure 15.**
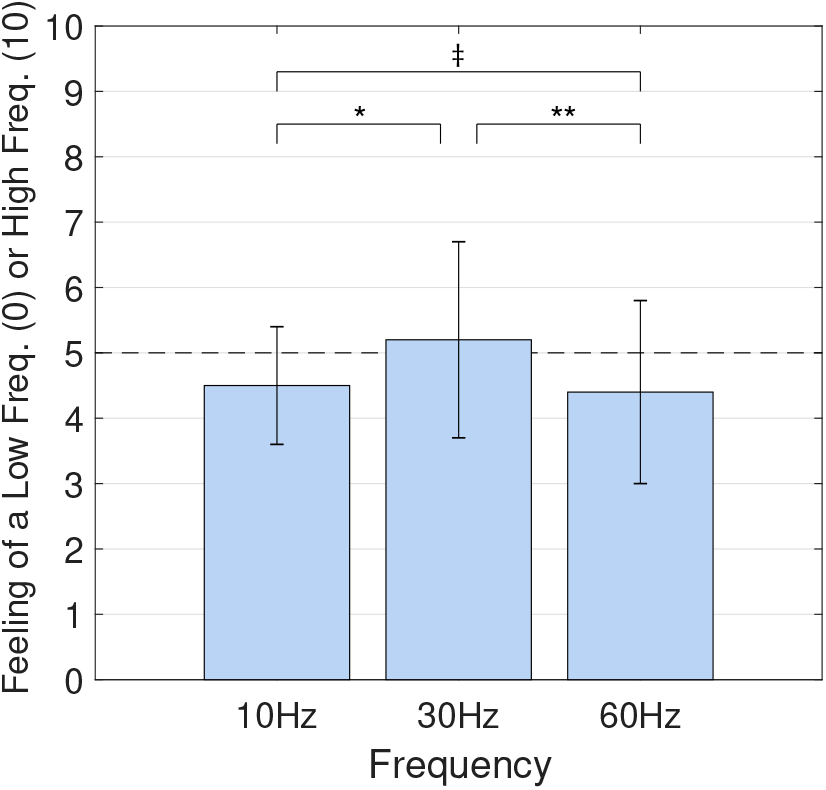
The effect of increasing signal frequency on the sensation of a low or high-frequency vibration is shown here. Significance of the overall differences between all groups: 0.523, \**p* = 0.367, * * *p* = 0.356, ‡*p* = 0.846

We can see in Fig. 15 that increasing the input signal frequency does not necessarily indicate an increase in the feeling of a higher frequency vibration. To the contrary, an input signal with a frequency of 30*Hz* induces a higher frequency vibration feeling than a 60*Hz* signal. The differences between the groups were not significant.

Furthermore, we can investigate the effect of duty cycle on this feeling too. Fig. 16 shows that this association is not notable. Both duty cycles tested in this study showed almost similar effect in relation to the feeling of a high or low-frequency vibration. The difference is not significant (*p* = 0.9).

**Figure 16.**
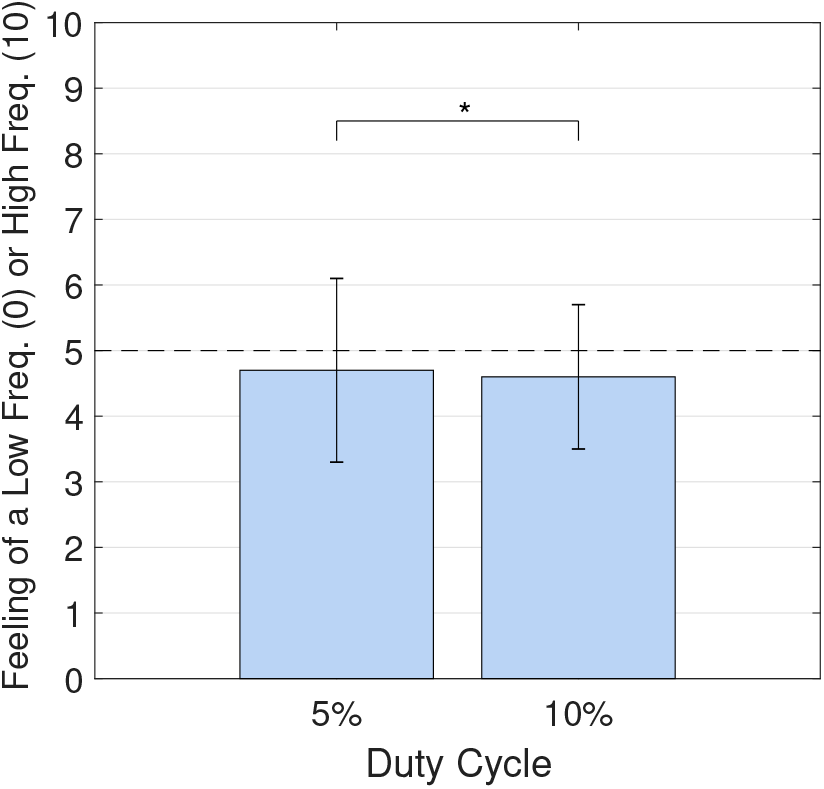
The effect of increasing signal duty cycle on the sensation of a low or high-frequency vibration is shown here. ∗*p* = 0.9

Finally, the impact of changing voltage can be studied too. This is shown in Fig. 17. The overall difference between groups and also the difference between each two groups are all statistically significant.

**Figure 17.**
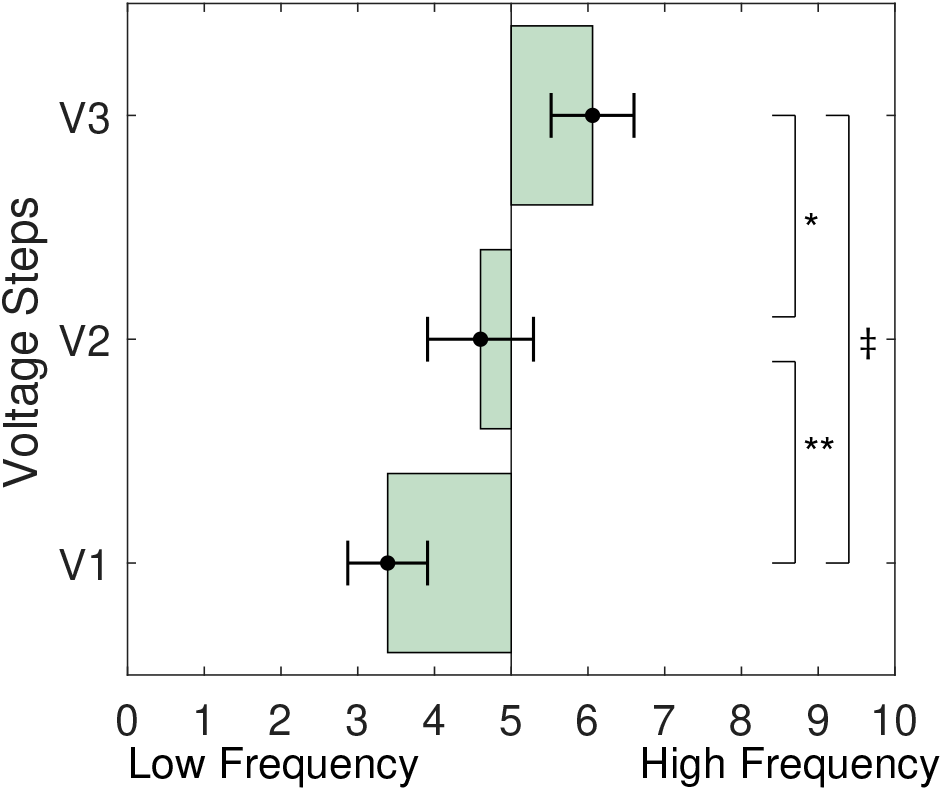
The effect of increasing signal voltage on the sensation of a low or high-frequency vibration is shown here. Significance of the overall differences between all groups: < 0.0001, ∗*p* = 0.003, ∗ ∗ *p* = 0.006, ‡*p* < 0.0001

A conclusion can be drawn here from Fig. 17 that voltage has the most pronounced effect on the feeling of a high or low-frequency in comparison with signal duty cycle or frequency. The lower voltage causes the experience of a feeling of low-frequency vibration but as we increased the voltage, the reported feelings are more of a high-frequency vibration. This can be explained by the fact that higher voltage caused the sensation to be intensified.

As an overall look to the relations between various signals and this sensation, a sorted plot is generated using all the 18 different signals that were tested in this study to see how they compare to each other in regard to the sensation of low or high vibration. Fig. 18 shows this arrangement. It can be seen that a signal with the frequency of 60*Hz* and duty cycle of 5% at the detection threshold voltage can give the lowest vibration feeling among all the signals tested. One fact to notice on this plot is how the standard deviation is almost equal for each of the tests and the average score has a steady increase. This shows that this sorting is very reliable based on the reports from the participants.

**Figure 18.**
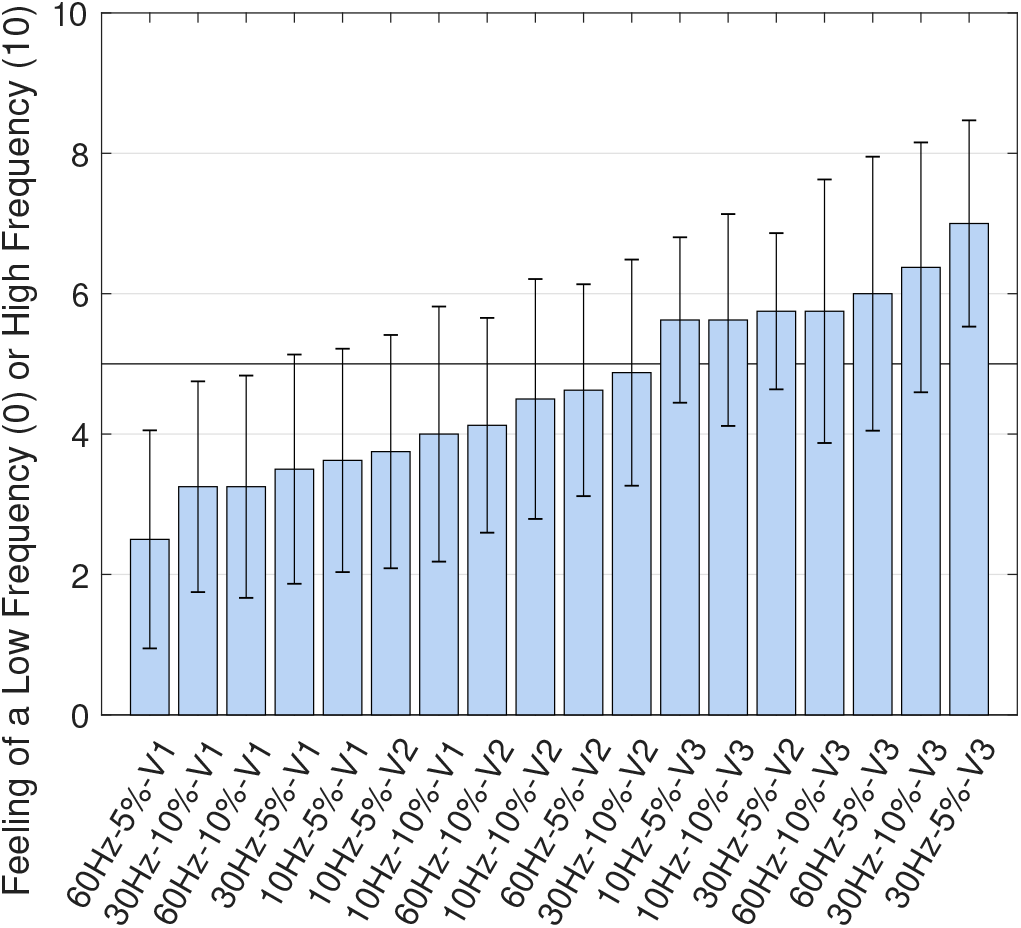
All the 18 different signals are sorted here based on the sensation of a low or high-frequency vibration.

#### 3) Surface and Shallow sensation vs. Deep Sensation

One of the feelings that were reported in previous studies and is a subject of interest, is whether the sensation is felt on the surface of the skin or it is felt deep inside the tissue. A surface sensation can also be expressed as a shallow sensation. To find the dominant factors that produce this feeling, each parameter of the input signal was examined.

Fig. 19 shows the relationship between the frequency of the signal and the sensation of a surface or deep sensation. Considering this figure, a noteworthy change does not exist as a function of changing the signal frequency. The overall differences between all groups is not significant. Although, it is known that higher frequencies would eventually cause a deeper sensation, but we can speculate that the difference between 30*Hz* and 60*Hz* is not large enough to show their effects here.

**Figure 19.**
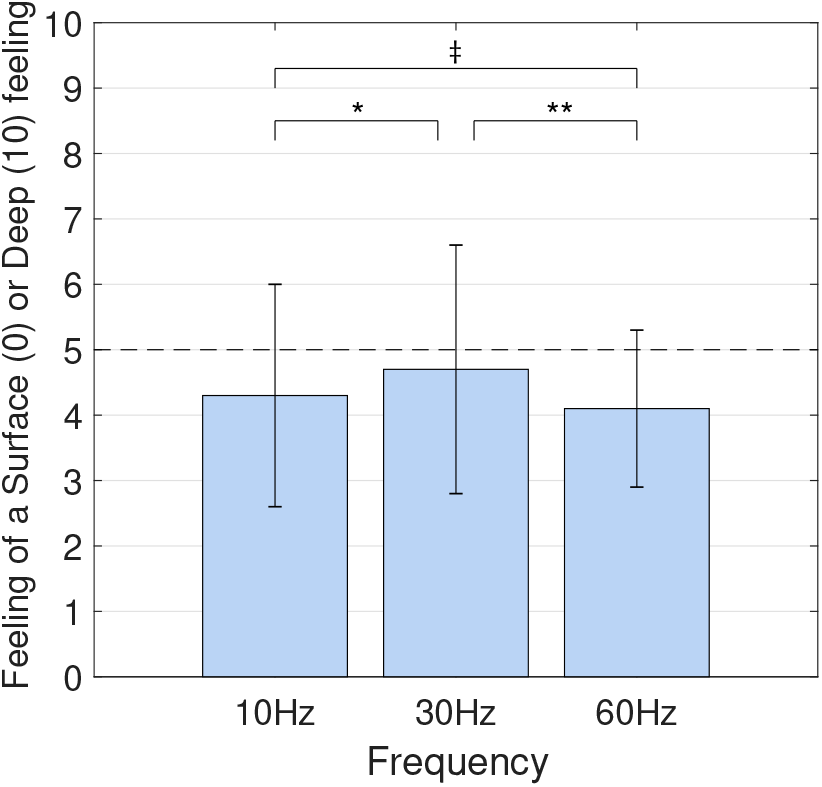
The effect of increasing signal frequency on the sensation of a surface or deep sensation is shown here. Significance of the overall differences between all groups: 0.822, ∗*p* = 0.702, ∗ ∗ *p* = 0.545, ‡*p* = 0.851

We can also investigate the effect of duty cycle on the changes of this feeling. Fig. 20 shows this relationship. Once again, the connection is not substantial and just a minute increase is seen as a result of a higher duty cycle. The difference is not significant (*p* = 0.6).

**Figure 20.**
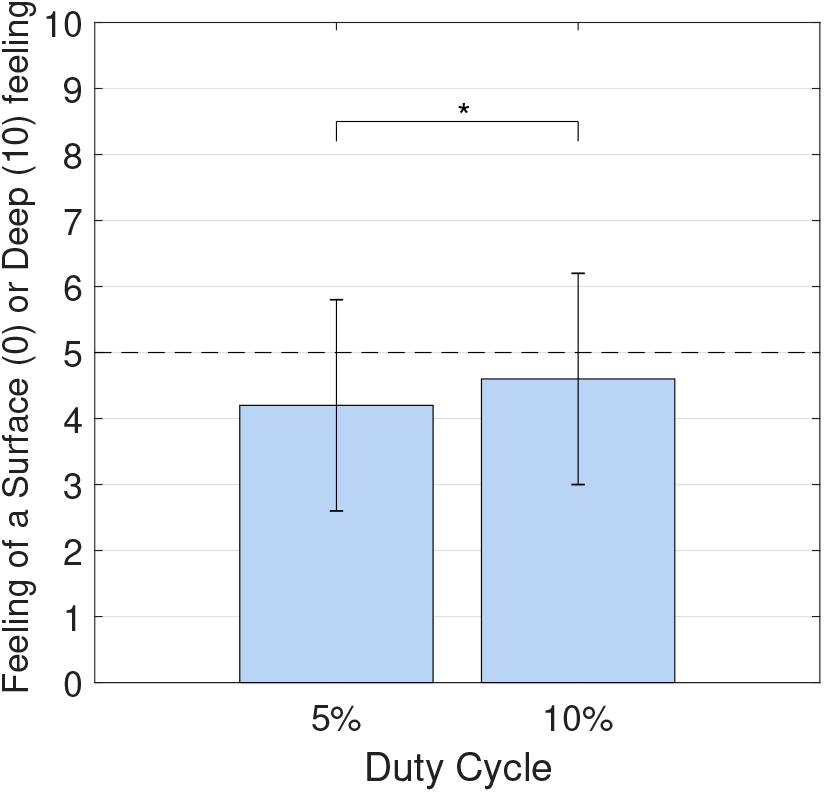
The effect of increasing signal duty cycle on the sensation of a surface or deep sensation is shown here. ∗*p* = 0.595

Finally, the effect of voltage can be inspected. Fig. 21 shows this relation. All of the differences here are statistically significant.

**Figure 21.**
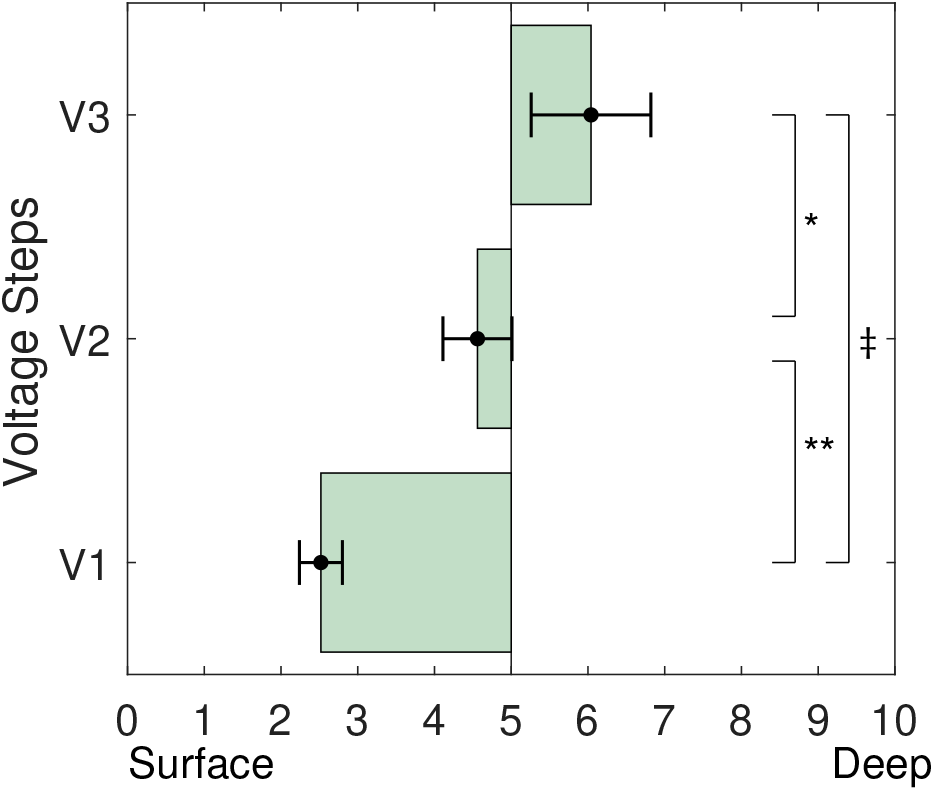
The effect of increasing signal voltage on the sensation of a surface or deep sensation is shown here. Significance of the overall differences between all groups: < 0.0001, ∗*p* = 0.003, ∗ ∗ *p* < 0.0001, ‡*p* < 0.0001

We can see one more time in Fig. 21 that changing the voltage is the ruling factor in causing a surface or deep sensation. This can be related to the more pronounced intensity that higher voltages cause.

To investigate which one of the 18 test signals better causes a surface or deep sensation, a sorted plot is generated and shown in Fig. 22; An almost perfect grouping, with just one exception, can be seen based on the voltage. Another noteworthy observation can be the gap or step between each voltage groups. It shows that increasing the voltage does in fact causes a definite and sudden shift toward deeper sensations.

**Figure 22.**
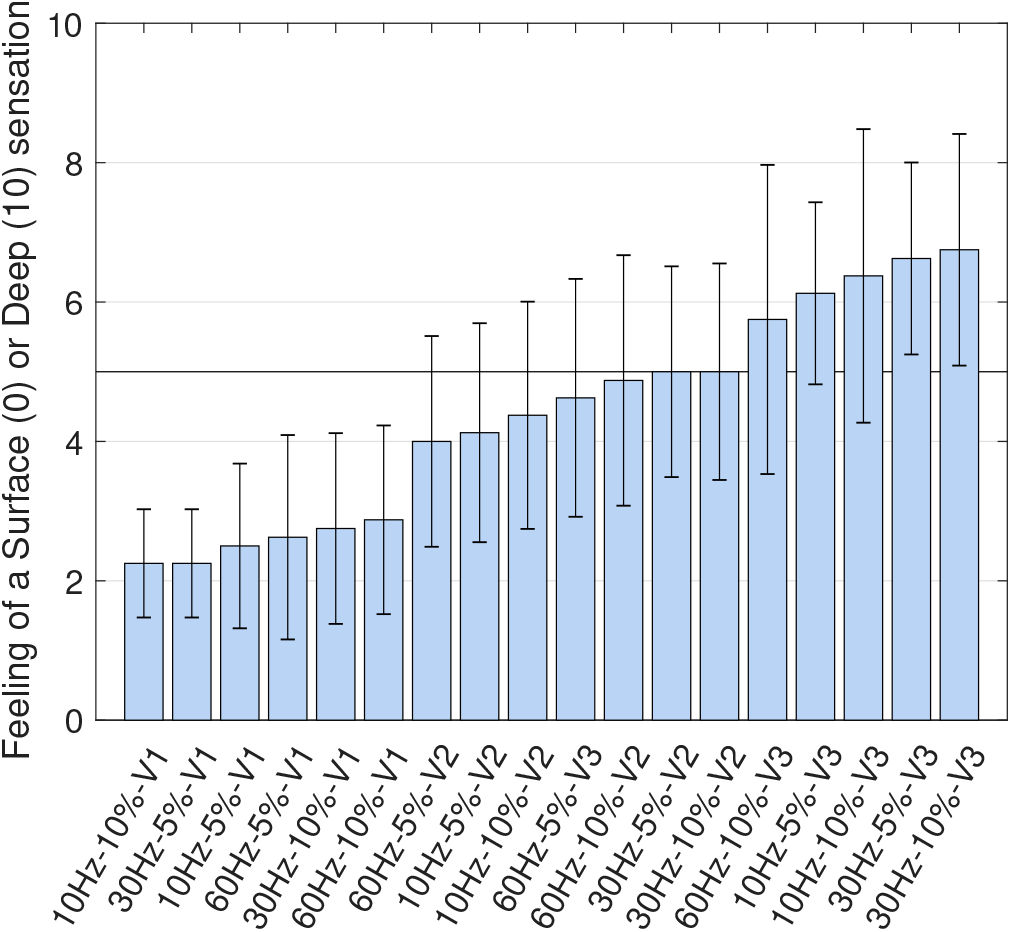
All of the 18 different signals are sorted here based on the sensation of a surface or deep sensation.

#### 4) Smooth vs. Rough

Three characteristics of the material sensations were investigated in this study. This can be later used to induce a specific material or object using a specially crafted electric signal. The goal is to see whether a special signal exists to produce the sensation of touching a specific surface material or not.

The first one of these material sensations was the feeling of a smooth or rough surface. It was explained to the participants whether they can interpret the signal as a smooth or rough surface, and has nothing to do with the contacts surface themselves. It was also described to them that this feeling could be translated to the sensation of touching a piece of leather versus sandpaper. Choosing number “5” in this question (as before) meant a “neutral” feeling indicating that the subject could not associate the signal with a specific material.

As with other sensations, each parameter of the input signal was examined individually to find the governing factors.

The effect of changes in duty cycle on the sensation of a smooth or rough surface is shown in Fig. 23. A small shift toward the smoothness can be seen when using a higher duty cycle but the difference is not significant (*p* = 0.34).

**Figure 23.**
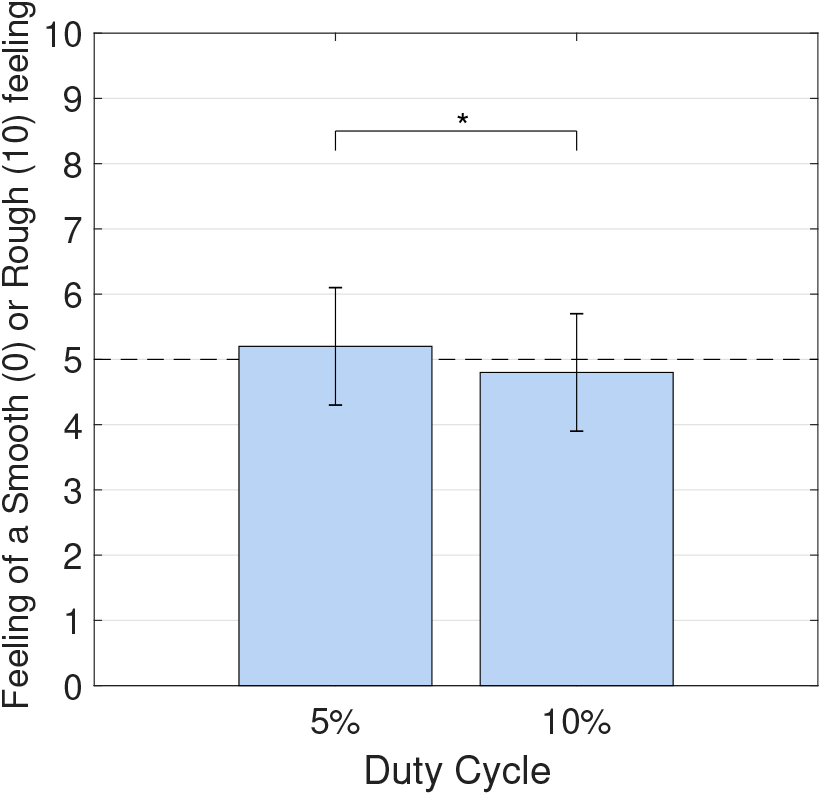
The effect of increasing signal duty cycle on the sensation of a smooth or rough surface is shown here. ∗*p* = 0.340

In the next step, the effect of changes in frequency was examined. Fig. 24 shows how the sensation of the surface was transferred to a smoother one by increasing the frequency. The small values of standard deviation show that this perception was prevalent across the participants. Still, the difference is not statistically significant.

**Figure 24.**
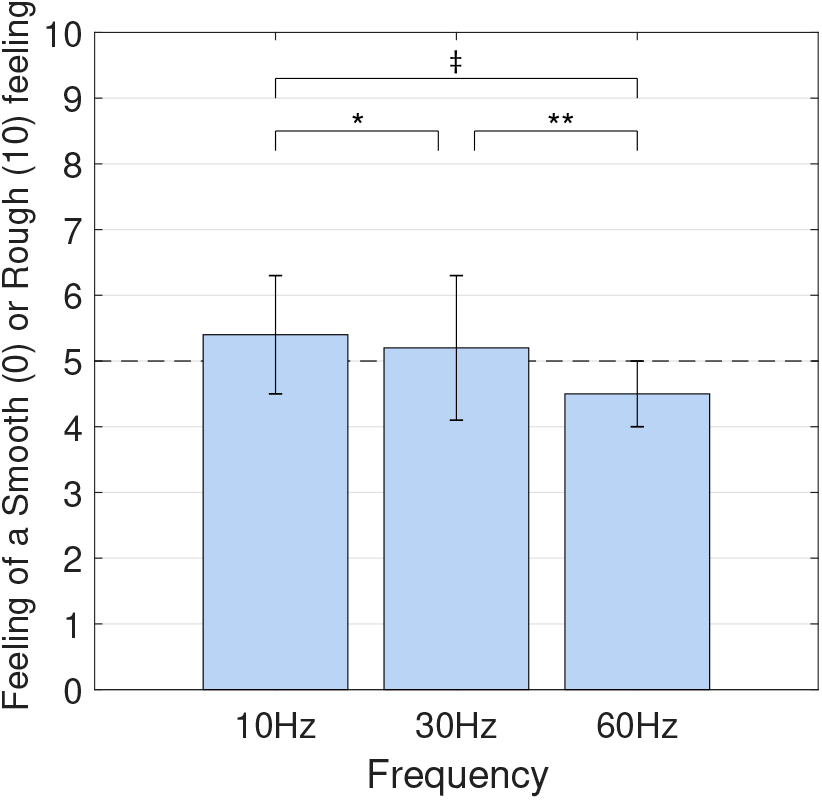
The effect of increasing signal frequency on the sensation of a smooth or rough surface is shown here. Significance of the overall differences between all groups: 0.247, ∗*p* = 0.735, ∗ ∗ *p* = 0.219, ‡*p* = 0.076

We were also interested in seeing the effects of voltage on the feeling of a smooth or rough surface. Fig. 25 shows this relationship. It can be seen that lower voltages induce a smoother surface whereas higher voltages give the perception of a rougher surface. The results are very symmetric in this regard as the middle voltage range falls exactly at “5” which means a neutral feeling. The differences between all the groups here are statistically significant.

**Figure 25.**
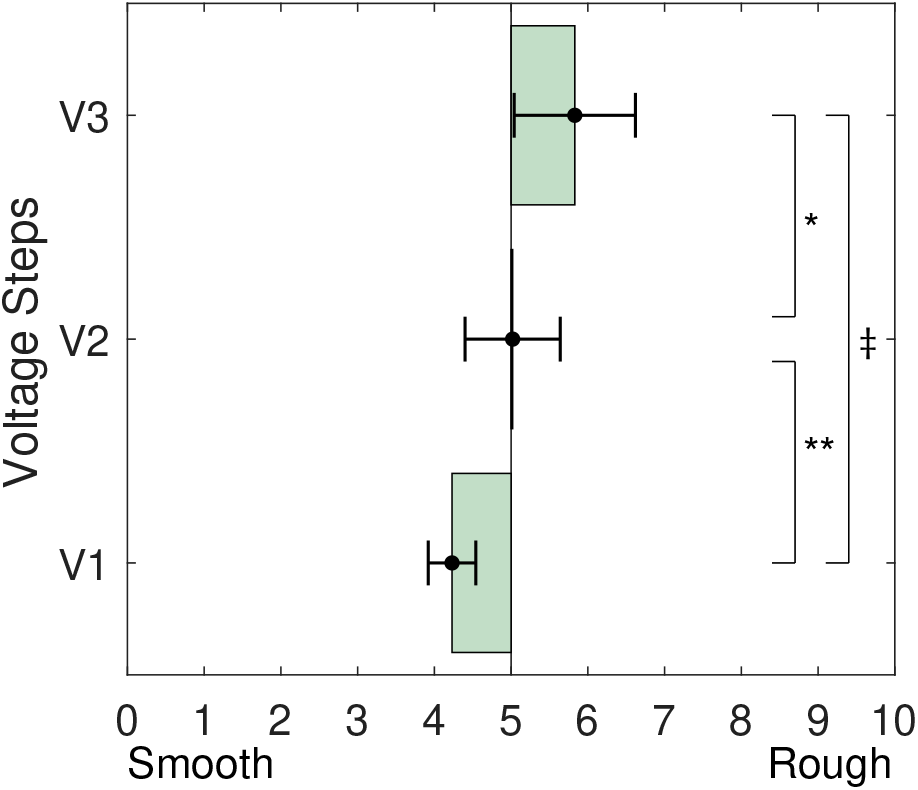
The effect of increasing signal voltage on the sensation of a smooth or rough surface is shown here. Significance of the overall differences between all groups: < 0.001, ∗*p* = 0.075, ∗ ∗ *p* = 0.018, ‡*p* < 0.001

All of the 18 different signals that were studied in this work can also be examined in one figure by sorting them based on the reports of sensations. Fig 26 shows the sorted plot of all the signals. The difference between the roughest signal and the smoothest one is not notable and the variation is small, but, we can say that the 30*Hz* signal with the duty cycle of 10% at the threshold detection voltage can result in the smoothest feeling.

**Figure 26.**
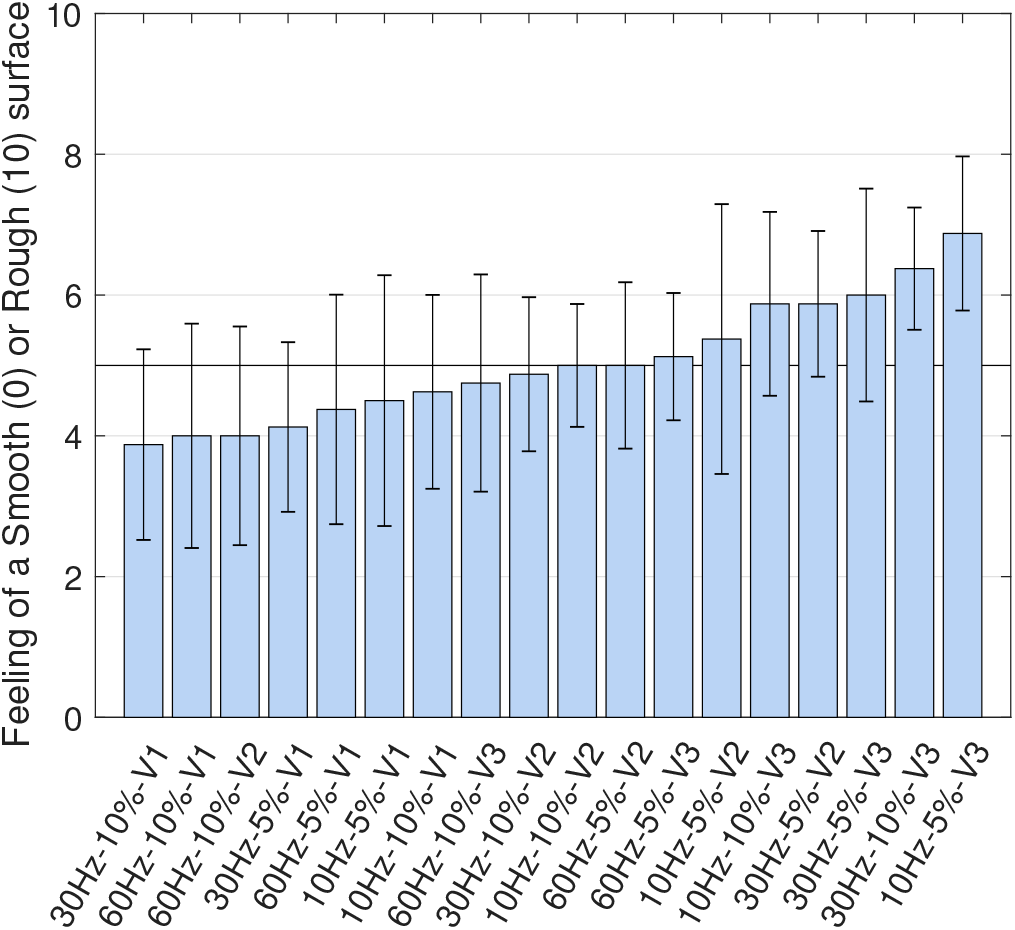
All the 18 different signals are sorted here based on the sensation of a smooth or rough surface.

#### 5) Flat vs. Bumpy

The second surface characteristic that was studied in this work was the sensation of a flat or bumpy surface. This can be translated to sensations of touching materials such as a tile or a sponge. Another example of a bumpy surface that was given to the participants was the surface of a keyboard versus the surface of a table.

The properties of the signal are examined individually here to find the effect of each property. First, we can see the result of changing the duty cycle in this sensation. Fig. 27 shows the association between the two. Noting the two groups, there is no differences in this regard (*p* = 0.9). We can conclude that changing the duty cycle is not a controlling factor in causing this sensation.

**Figure 27.**
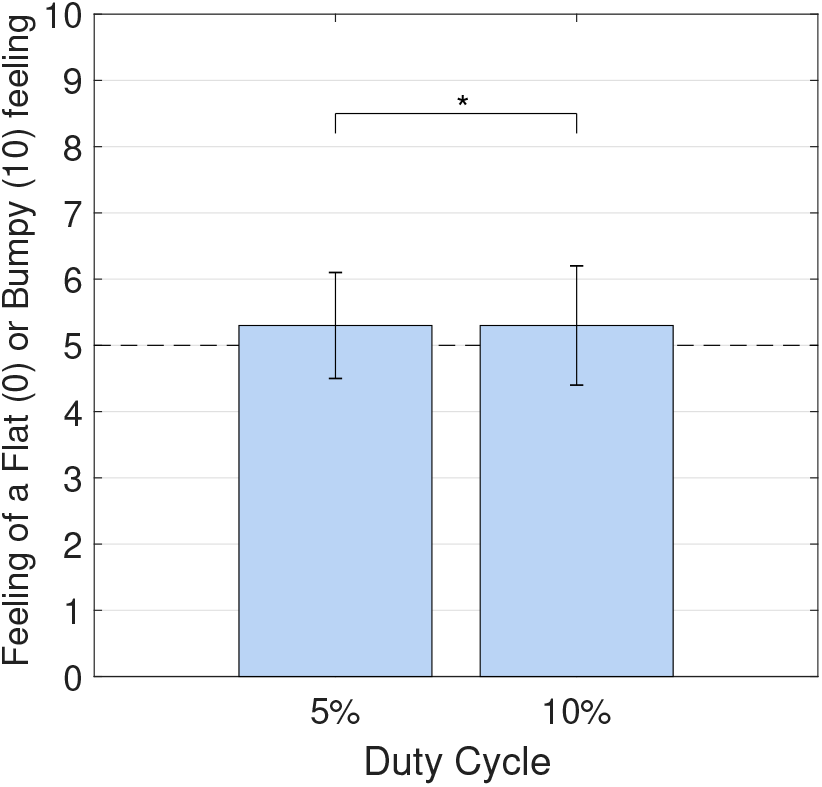
The effect of increasing signal duty cycle on the sensation of a flat or bumpy surface is shown here. ∗*p* = 0.872

Then, the effect of frequency on this sensation of the surface can be observed. Fig. 28 shows that higher frequencies are more likely to produce the sensation of a flat surface whereas a lower frequency can cause the feeling of a bumpy sensation. This might be the result of how Meissner corpuscles and Pacinian corpuscles are stimulated depending upon the frequency of the signal. The difference can be considered at 90% confidence level.

**Figure 28.**
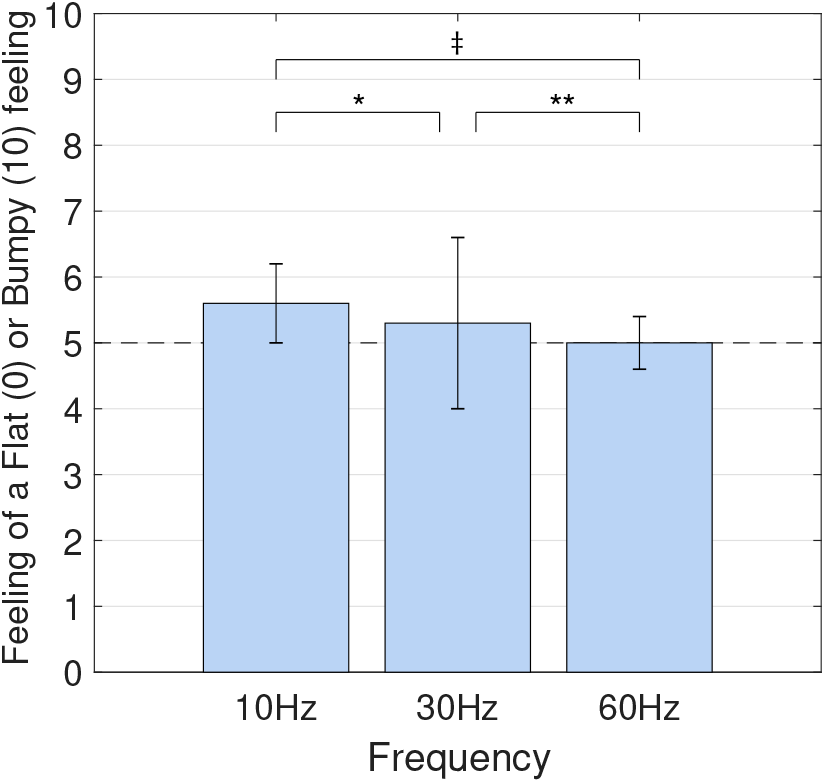
The effect of increasing signal frequency on the sensation of a flat or bumpy surface is shown here. Significance of the overall differences between all groups: 0.552, ∗*p* = 0.696, ∗ ∗ *p* = 0.583, ‡*p* = 0.1

Finally, the effect of voltage can be investigated too Fig. 29 indicated that once more, the voltage is playing the most important role in determining the sensation. Here the lower voltage of the signal resulted in a more flat surface sensation whereas any increases of the voltage can quickly cause a sensation of a bumpy surface. This is probably a result of stimulating the Meissner corpuscles. The differences between all the groups are statistically significant.

**Figure 29.**
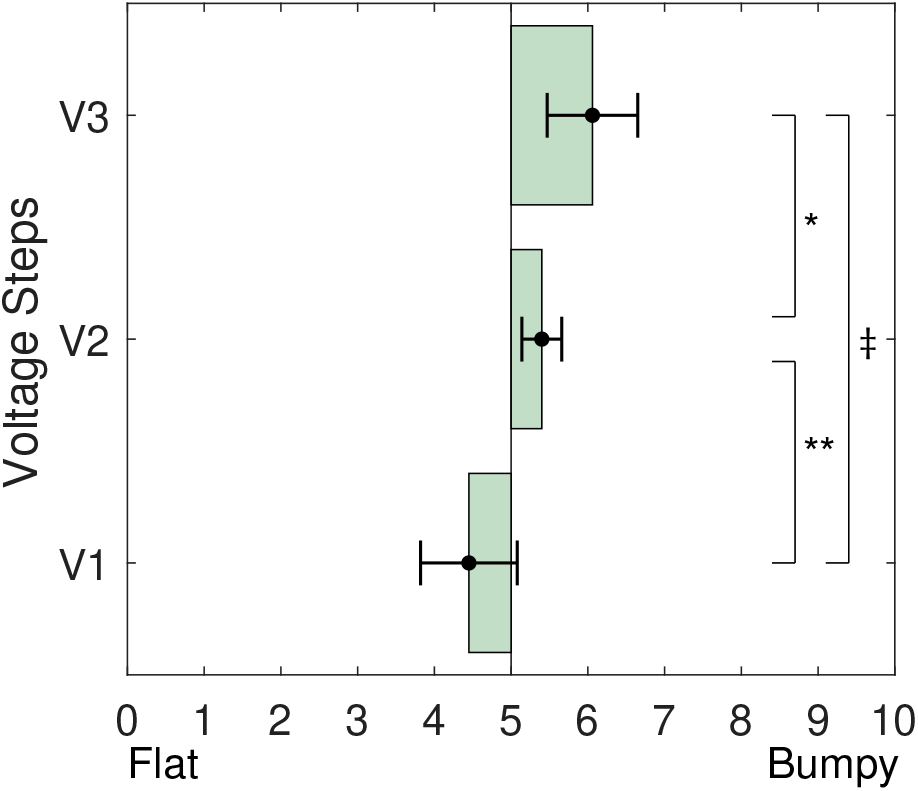
The effect of increasing signal voltage on the sensation of a flat or bumpy surface is shown here. Significance of the overall differences between all groups: < 0.001, ∗*p* = 0.030, ∗ ∗ *p* = 0.007, ‡*p* < 0.001

At the end, we can sort all of the 18 different signals based on the reports of a flat or bumpy surface sensation. Fig. 30 shows these arrangements. The range of variation is not significant. This can either show that the electrical stimulation is not a proper mean for inducing this sensation, or the range of signal properties that were studied in this work are not sufficient for this purpose. Based on this plot, the flattest sensation is caused by a signal with the frequency of 30*Hz* and the duty cycle of 5% at the detection threshold voltage.

**Figure 30.**
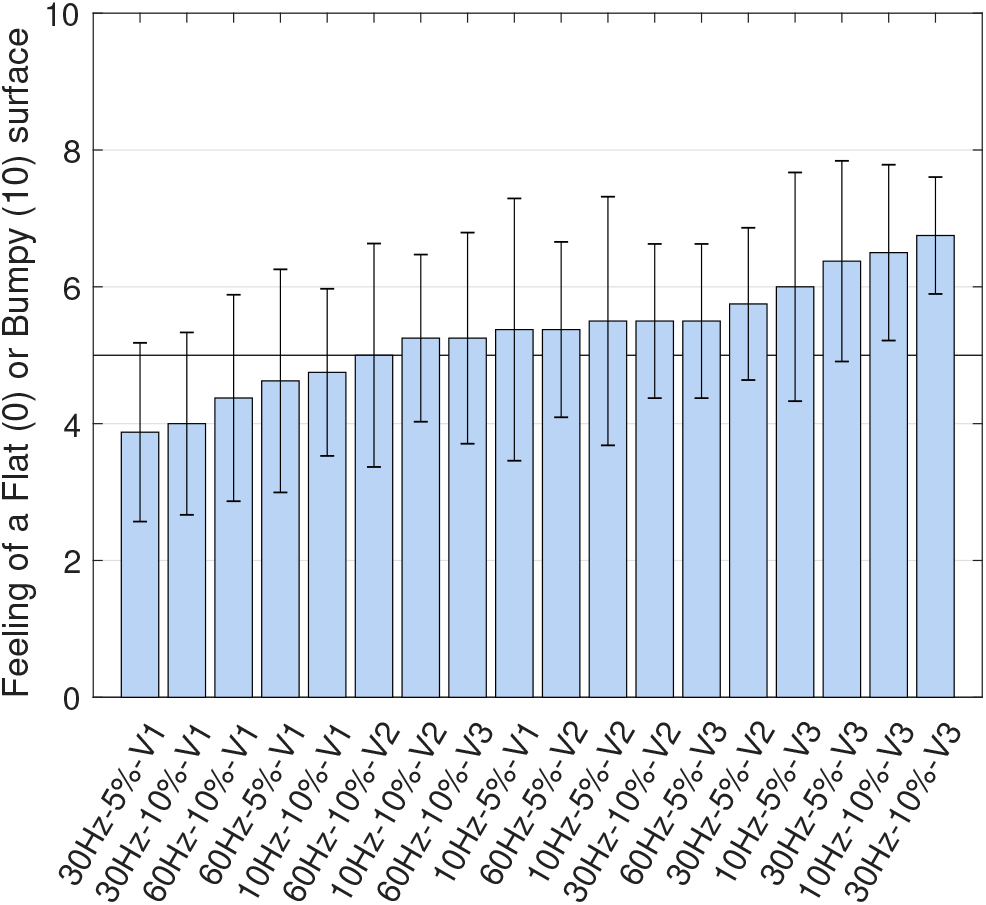
All the 18 different signals are sorted here based on the sensation of a flat or bumpy surface.

#### 6) Hard surface vs. Soft surface

The last surface characteristic that was studied in this work was the sensation of a soft or hard surface. Two examples were provided to the participants to facilitate understanding of the question. The first example was of a brick versus velvet which shows the hardness of the surface. The other example was the comparison between poking a table versus poking the surface of the hand where the muscles can flex as a result of the pressure.

The properties of the signals were inspected individually to discover the governing factors. Fig. 31 shows the effect of changing the duty cycle on this sensation. As it can be seen, changing the duty cycle does show an impact on the sensation of a hard or soft surface. Increasing the duty cycle can result in a sensation of a harder surface. The standard deviation is very small in this part. The difference between the two groups is statistically significant (*p* = 0.014). This shows that the duty cycle can be a controlling factor in regards to this sensation.

**Figure 31.**
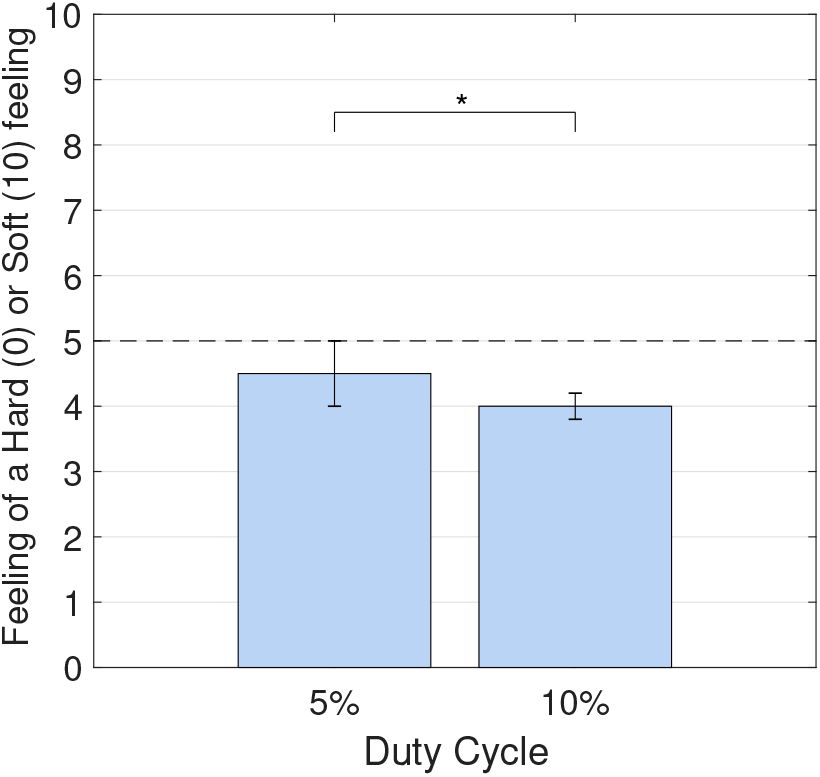
The effect of increasing signal duty cycle on the sensation of a hard or soft surface is shown here. ∗*p* = 0.014

The impact of changing the frequency can be examined too. Fig. 32 shows that frequency is not playing a major role in this regard. The overall tendency is toward a harder surface than a softer one. One possibility here is that the material of the contacts might have caused a deceptive impression on the participants as they may not have been able to report the sensation of the signal when their skin was actually touching the hard surface of the contacts. The differences between the groups are not significant.

**Figure 32.**
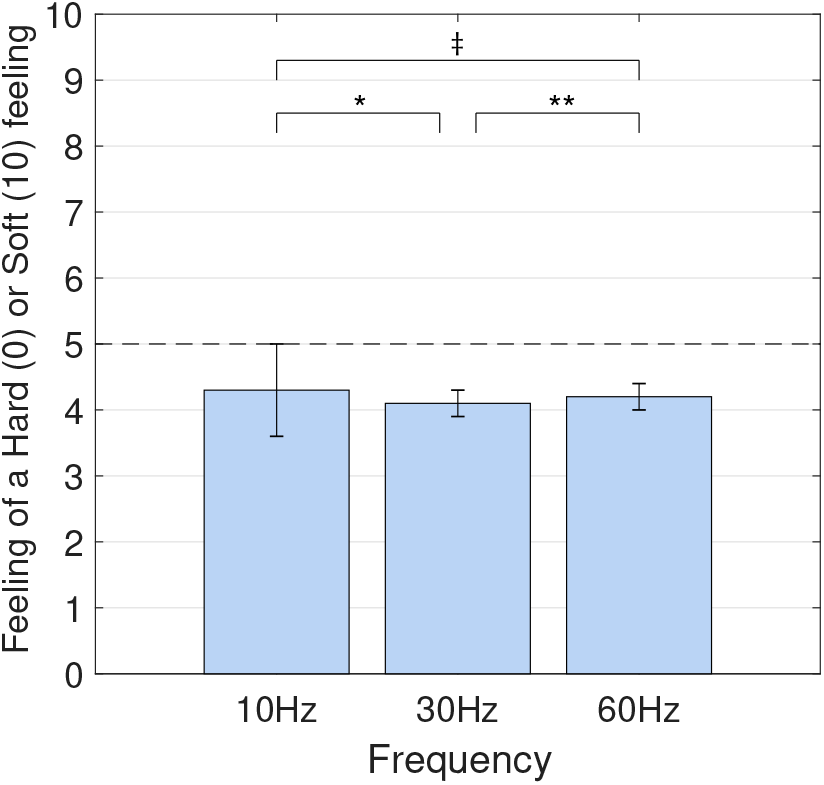
The effect of increasing signal frequency on the sensation of a hard or soft surface is shown here. Significance of the overall differences between all groups: 0.845, ∗*p* = 0.644, ∗ ∗ *p* = 0.417, ‡*p* = 0.918

We can also see the effect of changing the voltage on the perceiving sensation of a hard or soft surface. Fig. 33 shows that there are not significant differences as the result of changing the voltage (*p* = 0.8). For all the signals, the reports of the participants were leaning toward reporting a hard surface.

**Figure 33.**
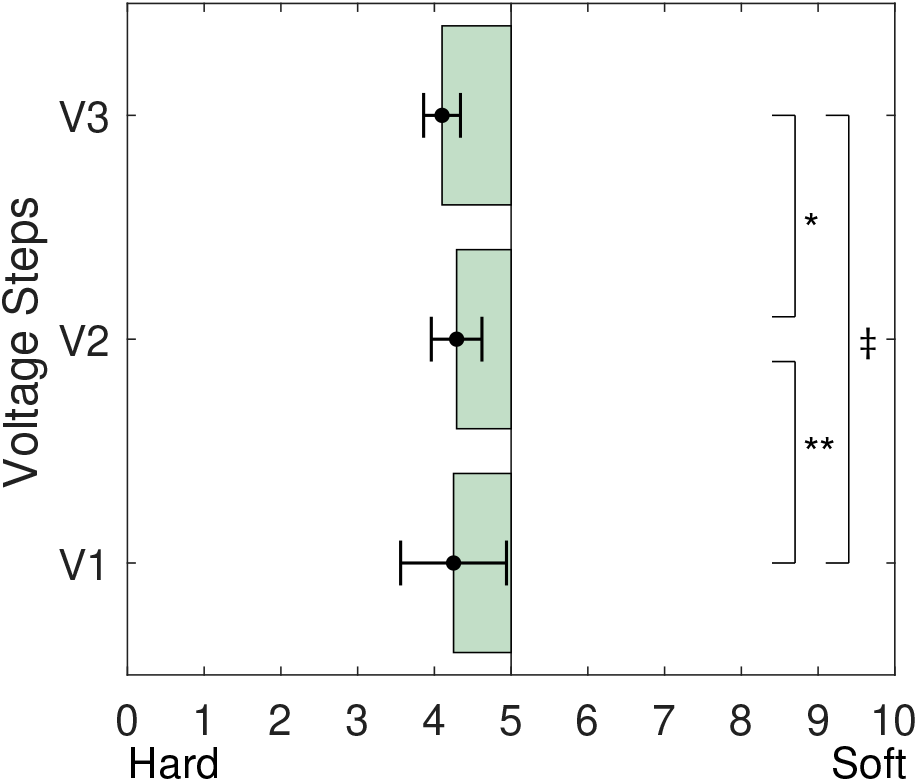
The effect of increasing signal voltage on the sensation of a hard or soft surface is shown here. Significance of the overall differences between all groups: 0.783, ∗*p* = 0.307, ∗ ∗ *p* = 0.877, ‡*p* = 0.664

At the end, all of the 18 signals can be investigated too. Fig. 34 shows a sorted plot of all the signals based on the reports of a hard or soft surface. The majority of the signals were reported as causing the sensation of a hard surface. As it was discussed before, this can either be a result of reporting the sensation of placing the skin on the contacts instead of the actual sensation of the signal, or it can be an indication that these ranges of variations in signal properties were not enough to show a significant differentiation in regard to the sensation of a hard or soft surface.

**Figure 34.**
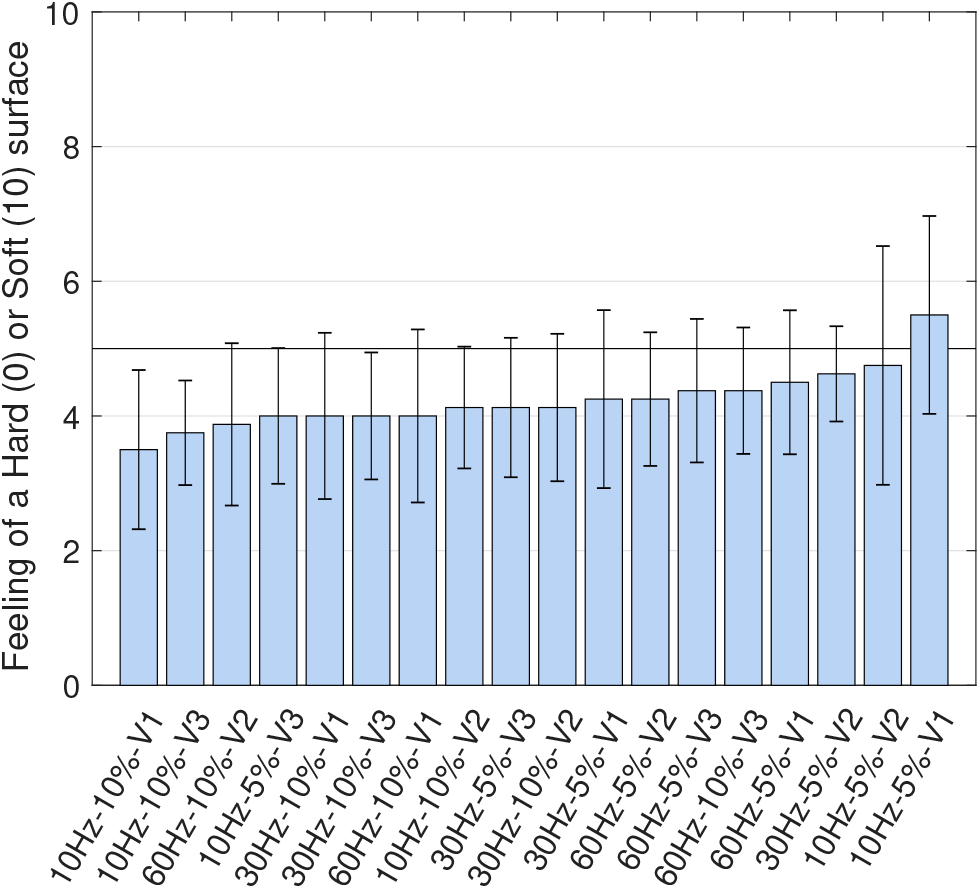
All the 18 different signals are sorted here based on the sensation of a hard or soft surface.

#### 7) Warm vs. Cold

Apart from the material properties, one factor that was investigated by other related research is the sensation of temperature. Although the same exact experiments were not done before but this is a topic of discussion. It is believed that it is possible to invoke a special electrical signal to stimulate a sensation of a warm or cold surface. Still, many of the participants in our study had a problem with associating this feeling with the sensations caused by the electrical signal.

The properties of the electric signal were investigated individually for this feeling. Fig. 35 shows how changing the duty cycle might affect the sensation of temperature. As it can be seen, both duty cycles stimulated a rather warm sensation but the difference between the two is not significant (*p* = 0.16).

**Figure 35.**
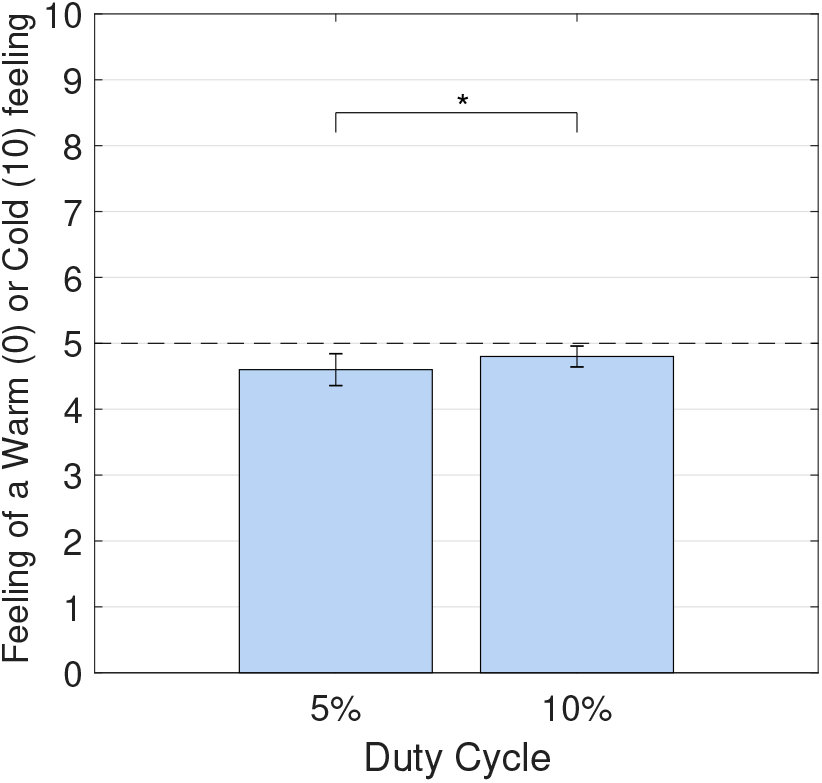
The effect of increasing signal duty cycle on the sensation of a warm or cold surface is shown here. ∗*p* = 0.160

The same examination can be done for the effect of changing the signal frequency. Fig. 36 shows this effect. Once again, it seems that this property does not affect the temperature sensation significantly (*p* = 0.347).

**Figure 36.**
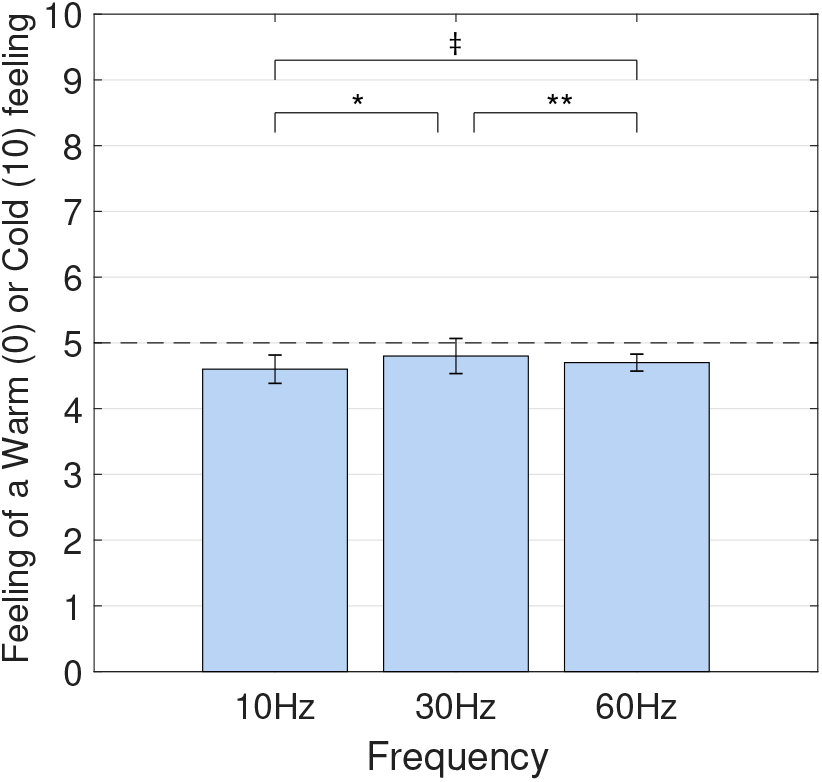
The effect of increasing signal frequency on the sensation of a warm or cold surface is shown here. Significance of the overall differences between all groups: 0.347, ∗*p* = 0.212, ∗ ∗ *p* = 0.591, ‡*p* = 0.313

Finally, the effect of changing the voltage can be investigated too. Fig. 37 shows this relationship. As was discussed before, most participants had a problem associating this feeling with the sensations of the electrical signal. Fig. 37 shows that most of the reports where around the neutral point with some leaning towards the warm sensation. The differences between the groups were not determined as being significant.

**Figure 37.**
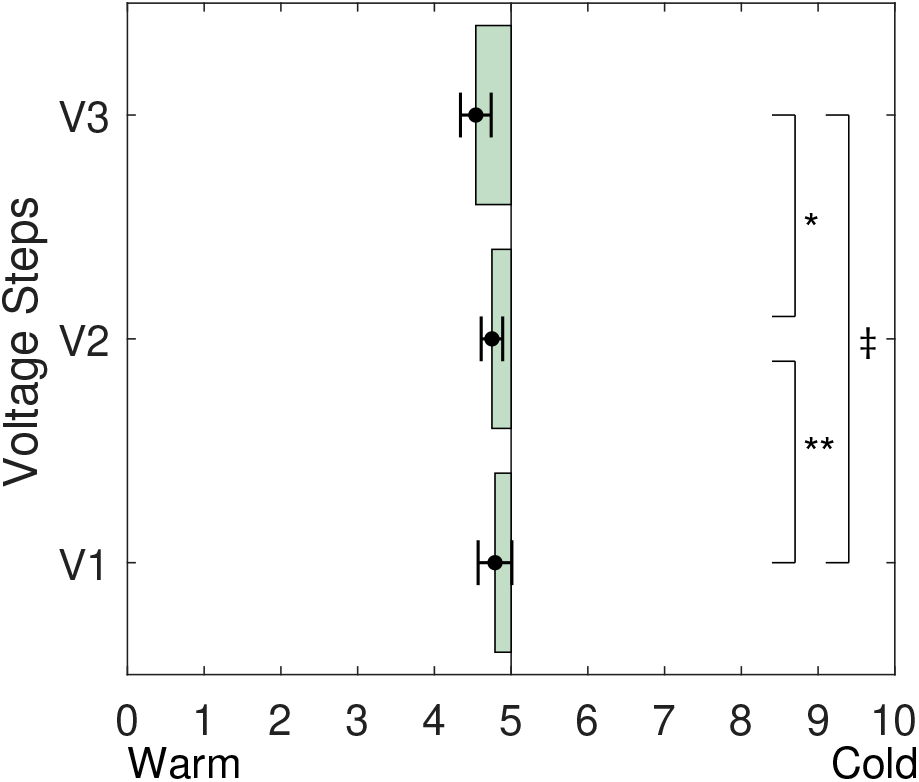
The effect of increasing signal voltage on the sensation of a warm or cold surface is shown here. Significance of the overall differences between all groups: 0.054, ∗*p* = 0.043, ∗ ∗ *p* = 0.760, ‡*p* = 0.049

At the end, all of the tested signals can be sorted based on the reports that were given on this sensation. Fig. 38 shows this sorted plot. Although all of the signals were interpreted as giving a warm sensation, there are very close to the neutral point (5). We can see that the participants were not able to properly associate this sensation with any of the signals provided in the experiments.

**Figure 38.**
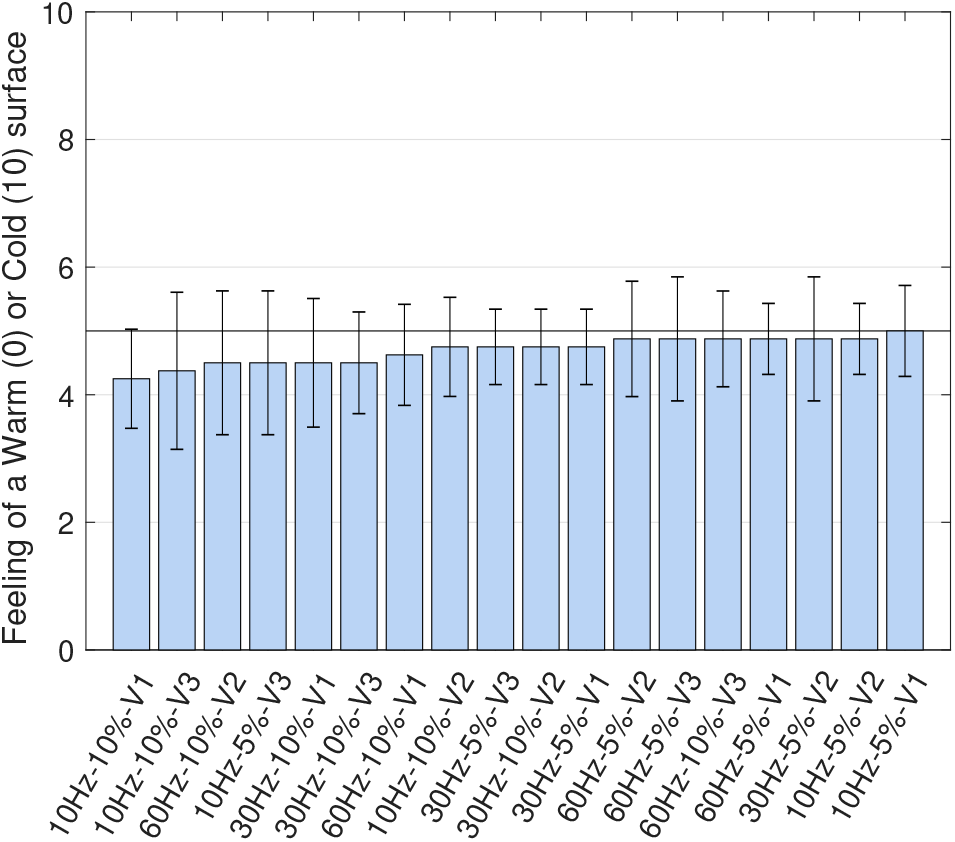
All the 18 different signals are sorted here based on the sensation of a warm or cold surface.

### C. Multidimensional Scaling Analysis

In this study, we have examined 18 different signals with various properties. It can be beneficial to categorize these signals to find out how and based on which characteristics do they relate to each other and form groups.

Multidimensional Scaling (MDS) analysis can be used to organize these signals into perceptual space where the distance between two objects shows their perceptual differences [21] [22]. A three-dimensional space is formed to present a satisfactory representation of the data. For adjusting the stress, the Kruskal’s stress method was used. The proximity matrix (using Euclidean distance) is shown in Table I.

**Table I.**
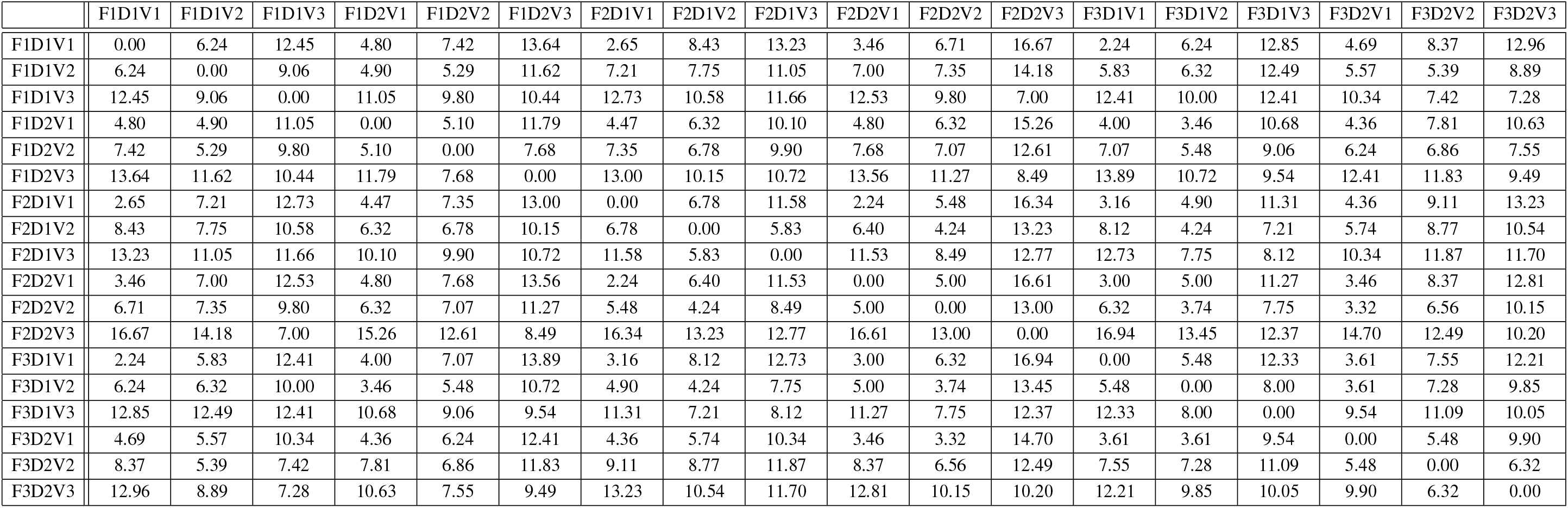
The proximity matrix (using Euclidean distance) for the Multidimensional Scaling analysis

The simple interpretation of the MDS is that as the similarity between the objects increases, the distance between them in the plot would decrease. This gives us a simple visual representation of how these signals are related and which characteristics can better bond or differentiate these signals.

The result of the MDS is shown in Fig. 39. This is a 3D representation of the analysis and each signal is shown as a ball in the space. The distance between them is an indication of how similar they are to one another. Three brackets were added manually to the plot to emphasise the grouping that is formed and found by MDS. Looking at this plot, it is noteworthy to see how these signals are grouped based on their voltage rather than any of the other properties. This shows that voltage is the best differentiating characteristic of these signals.

**Figure 39.**
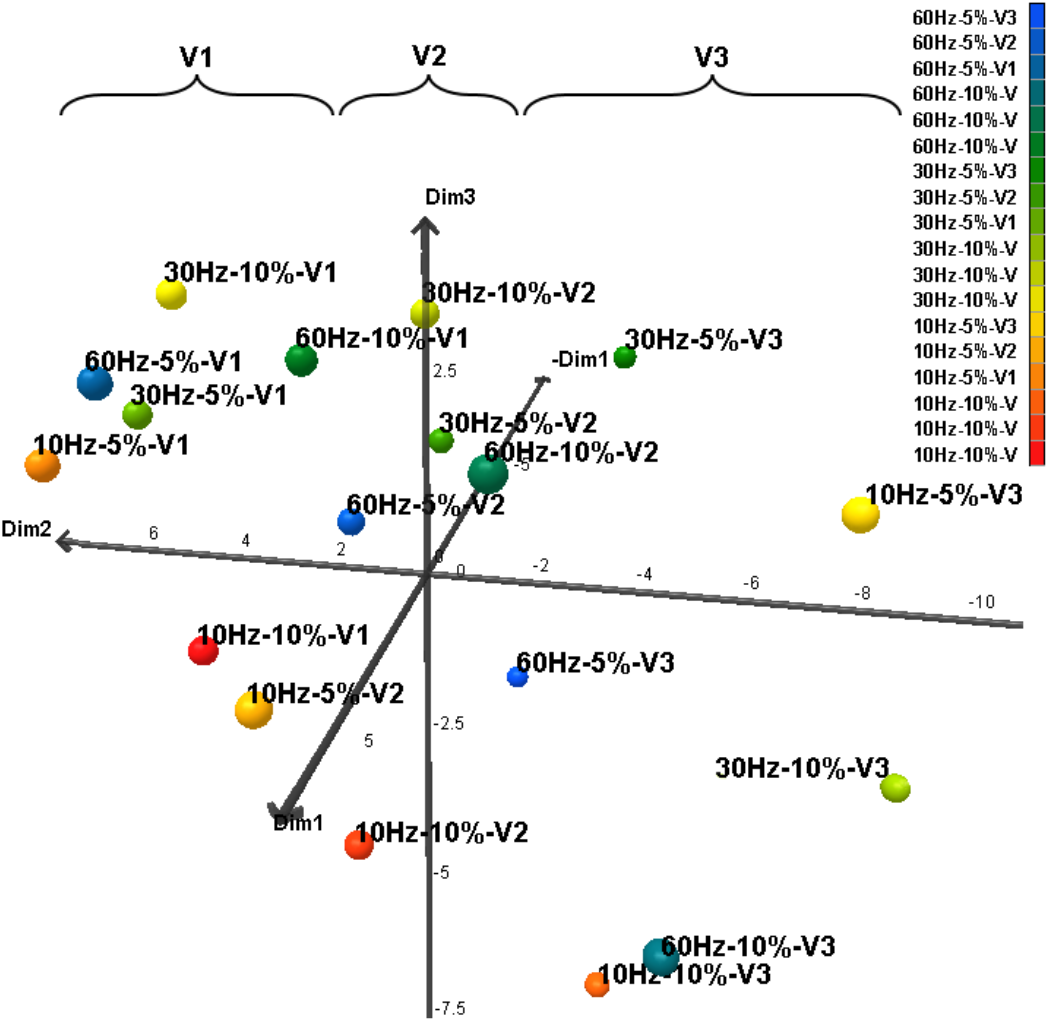
All the 18 different signals are analysed here using Multidimensional Scaling analysis. The brackets at the top are added manually to emphasis the grouping that was recognized by MDS. This figure shows that the most important and differentiating characteristic of all these signals is the voltage they are operating at. This shows that the frequency and duty cycle of the signals are not the governing factors in differentiating them.

## IV. Discussion

This study aims to determine the properties of the electrical signal that can result in a specific sensation. The results from the first part of the questionnaire can be summarized in a few facts. One realization is that the most conspicuous sensation that was felt by the participants was the feeling of “Vibration”. Meanwhile, the second most important feeling to them was the sensation of the “Slight touch”. Some of the sensations did not show a significant presence in the subjects’ reports. These were the sensations of “Unpleasant”, “Sting”, “Pressure” and “Deep Pressure”.

A pronounced trend was seen in the box plots that showed how the sensation of the “Slight touch” and “Sting or Unpleasant” was directly related to the voltage of the signal. It was shown that at the lowest voltage, the sensation of “Sting or Unpleasant” was very weak whereas when we increased the voltage, this sensation had strengthened.

We also saw the trends in all the descriptive characteristics versus the changes in each of the parameters of the signal. After analyzing the trends, in regards to the intensity of these sensations, it was determined that frequency had a slight influence on the sensation whereas the voltage affected many of them significantly. The duty cycle did not effect them at all.

In analyzing the second part of the questionnaire, each of the signal properties were examined individually to link them with the sensations. The signals were also examined collectively in sorted plots to see their associations to the sensation properties. One of the most important results that was seen in almost all of these analyses was how voltage had the strongest effect on the reported sensation rather than other properties. The frequency and duty cycle were analyzed too, but they were seldom a controlling factor. The p-values for all the tests of significance of differences are shown in Table II.

**Table II.** The p-Values for the test of significance across all the variables. (95% Confidence Level)

| p-values | Freq. (overall) | 10Hz / 30Hz | 20Hz / 60Hz | 10Hz / 60Hz | Voltage (overall) | V2 / V3 | V1 / V2 | V1 / V3 | Duty Cycle |
| --- | --- | --- | --- | --- | --- | --- | --- | --- | --- |
| Comfort / Uncomfortable | 0.930 | 0.988 | 0.760 | 0.707 | **<0.0001** | **<0.001** | **<0.0001** | **<0.0001** | 0.481 |
| Low freq. / High freq. feeling | 0.523 | 0.367 | 0.356 | 0.846 | **<0.0001** | **0.003** | **0.006** | **<0.0001** | 0.900 |
| Surface / Deep sensation | 0.822 | 0.702 | 0.545 | 0.851 | **<0.0001** | **0.003** | **<0.0001** | **<0.0001** | 0.595 |
| Smooth / Rough | 0.247 | 0.735 | 0.219 | 0.076 | **0.001** | 0.075 | **0.018** | **<0.001** | 0.340 |
| Flat / Bumpy | 0.552 | 0.696 | 0.583 | 0.100 | **<0.001** | **0.030** | **0.007** | **<0.001** | 0.872 |
| Hard / Soft | 0.845 | 0.644 | 0.417 | 0.918 | 0.783 | 0.307 | 0.877 | 0.664 | **0.014** |
| Warm / Cold | 0.347 | 0.212 | 0.591 | 0.313 | 0.054 | **0.043** | 0.760 | **0.049** | 0.160 |

The importance of the voltage level was also shown in the sorted plot of the “Comfortable” versus “Uncomfortable” sensation. The perfect grouping based on the voltage and also the steps in relation to the voltage groups showed how significant this factor is. The same was seen in the analysis of a low-frequency vibration feeling versus a high-frequency vibration feeling. Voltage had a significant effect there too. This was also true for the sensation of “Surface” versus the “Deep” sensation to some extent but the frequency was playing a slight role there nonetheless.

The characteristics of the material sensation were hardly expressed through the changes in signal frequencies.

One observation could be that low voltage of the signal is indirectly connected to “good” feelings such as “Comfortable”, “Low-frequency vibration” or “Smooth surface”. This might be because of how the participants have interpreted each parameter unconsciously. Based on these observations, it is clear that the low voltage signal was more desirable to the participants.

The Multidimensional Scaling analysis was the final conclusion to the previous reasonings in this work. MDS helped to show how these signals with various properties could be categorized and grouped together. The final conclusion was drawn once again that voltage is the differentiating factor. Noting Fig. 39, it is obvious that signals with different frequencies can be scattered across the defined space. The same was true about the duty cycle but this could not be said about the voltage. This was shown in the grouping that had occurred based on having the same voltage.

A question remains about how contradictory sensations can be achieved. For example, how can we have a signal that is producing a comfortable sensation, but at the same time, gives the sensation of a rough surface. This question is probably rooted in the paradox of the perceptual processes that are involved as the side effects of these experiments. We might not be able to produce such signals that have characteristic with dependence on conflicting properties of the input signal. In other words, we know that low voltage is more comfortable and we also know that to stimulate a rough sensation, we need a high voltage. This conflict cannot be resolved solely by adjusting the voltage. Further investigations are needed to find out how such behaviors could be generated.

## V. Conclusion

This study examined the relationship between various signal properties and the resulting feelings induced on the skin by conducting 144 measurements. The properties studied included voltage, frequency, and duty cycle. A total of 18 signals were composed by combining several options of these properties. A questionnaire was designed to examine the reports of the participants. In this study, 8 participants contributed with their reports on different signals. The results were analyzed using various statistical methods.

One overall conclusion was that most participants felt a feeling of “Vibration” as the result of applying the electrical signal to their skin. Most signals were also reported as being “Comfortable”.

It was shown that increasing the voltage, rather than any of the other parameters, was responsible for causing an “Uncomfortable” feeling.

In another part of the study, the relationship between each of the signals and different categories of feelings was studied.

It was shown that a feeling of “Low-frequency” vibration can be better produced with lower voltages and does not necessarily depend on the frequency of the electro-tactile signal itself. The same could be also said about creating a sensation on the “Surface” of the skin versus a “Deep” sensation. Here again, the voltage was the governing factor and the frequency had limited effects.

Some characteristics of the material were investigated as well.

Regarding the sensation of a “Smooth” or “Rough” surface, it was shown that lower voltages can lead to a feeling of a “Smooth” surface whereas a higher voltage was inducing a “rougher” surface. In this part, the higher frequency was contributing to a sensation of a “smoother” surface.

The flatness of the surface was studied as one of the characteristics of the surface material. The effect of the voltage was less significant in this part, but still, a lower voltage was related to a more “flat” surface when a higher voltage was associated with a “bumpy” surface. Frequency had a minute effect in this regard but higher frequencies were more related to a feeling of a “flat” surface.

The final characteristic of the surface studies here was the “softness”. Almost all of the signals were perceived by the participants as a hard surface and we could not associate any signals with a soft surface. Also, frequency did not play a role here. On the other hand, this was the only category that was shown to be influenced by the duty cycle. The bigger duty cycle did induce the feeling of a harder surface.

Finally, the sensation of a warm or cold surface was also examined. The participants could not associate the electrical signal with any of these feelings and most of the reports were “neutral”.

Duty cycle did not play a role in most of these experiments; In some of the tests that duty cycle had a possibility of an effect, it was not significant.

To better examine the governing factors of these various signals, a Multidimensional Scaling analysis was conducted and the results showed that among voltage, frequency, and duty cycle, the most differentiating and governing role is played by the voltage. It was seen that all of the 18 signals were grouped based on their voltages rather than any of the other properties.

## Notes

The research work was partially supported by NSF CAREER Award CBET#1352006.

### Competing Interest Statement

The authors have declared no competing interest.

